# The ER cholesterol sensor SCAP promotes CARTS biogenesis at ER-Golgi contact sites

**DOI:** 10.1101/679936

**Authors:** Yuichi Wakana, Kaito Hayashi, Takumi Nemoto, Chiaki Watanabe, Masato Taoka, Felix Campelo, Hidetoshi Kumata, Tomonari Umemura, Hiroki Inoue, Kohei Arasaki, Mitsuo Tagaya

**Author notes:** Corresponding authors: Yuichi Wakana and Mitsuo Tagaya. K. Hayashi and T. Nemoto contributed equally to this paper.

## Abstract

In response to cholesterol deprivation, SCAP escorts SREBP transcription factors from the endoplasmic reticulum (ER) to the Golgi complex for their proteolytic activation, leading to gene expression for cholesterol synthesis and uptake. Here we show that in cholesterol-fed cells ER-localized SCAP interacts through Sac1 phosphoinositide 4-phosphate (PI4P) phosphatase with a VAP/OSBP complex, which mediates counter-transport of ER cholesterol and Golgi PI4P at ER-Golgi contact sites. SCAP knockdown inhibited the turnover of PI4P perhaps due to a cholesterol transport defect and altered the subcellular distribution of the VAP/OSBP complex. As in the case of perturbation of lipid transfer complexes at ER-Golgi contact sites, SCAP knockdown inhibited the biogenesis of the *trans*-Golgi network-derived transport carriers CARTS, which was reversed by expression of wild-type SCAP but not cholesterol sensing-defective mutants. Altogether, our findings reveal a new role of SCAP under cholesterol-fed conditions in the facilitation of CARTS biogenesis at ER-Golgi contact sites, depending on the ER cholesterol.

**Summary:** SCAP is the key regulatory protein in cholesterol metabolism. Wakana et al. describe a new role of SCAP in controlling Golgi PI4P turnover and the biogenesis of the Golgi-derived transport carries CARTS via cholesterol/PI4P exchange machinery at ER-Golgi contact sites.

## Introduction

Cholesterol plays a crucial role in regulating the functions of mammalian cell membranes by determining their integrity and fluidity. Cholesterol together with sphingolipids can form liquid-ordered membrane nanodomains which are segregated from other lipids and thus are proposed to serve as platforms for specific proteins that regulate signal transduction and endocytosis at the plasma membrane (PM), and apical transport from the *trans*-Golgi network (TGN)^1–8^. Increasing evidence –including recent experiments where sphingomyelin (SM) metabolism at the *trans*-Golgi membranes was perturbed– strongly suggests that such lipid nanodomains are required for the functional organization of enzymatic domains, cargo sorting, and transport carrier biogenesis at the TGN^9–14^, and thereby are crucial to maintain homeostatic control of the Golgi function.

*De novo* biosynthesis of cholesterol occurs at the endoplasmic reticulum (ER), where the key regulatory protein in cholesterol metabolism, called sterol regulatory element binding protein (SREBP) cleavage-activating protein (SCAP), localizes. SCAP is a polytopic membrane protein that functions as an ER cholesterol sensor to control the cellular cholesterol content^15–20^. When the ER cholesterol level is low, SCAP escorts the membrane-bound transcription factors SREBPs into COPII vesicles for their export from the ER to the Golgi complex. There, SREBPs are cleaved by proteases, allowing their transcriptionally active domain to enter the nucleus and promote expression of genes involved in cholesterol synthesis and uptake. Conversely, when cholesterol in the ER membrane is abundant, cholesterol binds to SCAP and triggers conformational changes that allow SCAP to interact with the integral ER membrane protein Insig, which retains SCAP in the ER along with unprocessed SREBPs. However, it remains unclear whether SCAP has other functions under cholesterol-fed conditions, except for sequestering SREBPs at the ER.

Although the ER produces cholesterol and also receives it from the PM and other sources, ER cholesterol content is low^21^. This is accomplished through the export of cholesterol from the ER against the concentration gradient in both vesicular and non-vesicular manners, the latter of which occurs at membrane contact sites. In particular, at ER-Golgi contact sites, Golgi-associated oxysterol-binding protein (OSBP) interacts with an integral ER membrane protein named vesicle-associated membrane protein– associated protein (VAP), to transfer cholesterol from the ER to the *trans*-Golgi membranes, accompanied by reciprocal transfer of phosphoinositide 4-phosphate (PI4P)^22^. PI4P transported to the ER is hydrolyzed by the ER-localized lipid phosphatase Sac1, which thus seems to provide a driving force for cholesterol transport^23^.

ER-Golgi contact sites also control the transport of ceramide from the ER to the *trans*-Golgi membranes. This non-vesicular transport is mediated by a complex of VAP and ceramide transfer protein (CERT), and leads to the biosynthesis of SM and diacylglycerol (DAG) by SM synthase at the *trans*-Golgi membranes^24–26^. Importantly, PI4P in the *trans*-Golgi membranes is crucial for both OSBP and CERT to associate with the Golgi complex^27^ and hence the biosynthesis and metabolism of cholesterol, SM, PI4P, and DAG are either directly or indirectly interconnected^28^. Of note, DAG recruits to the TGN a serine/threonine kinase, protein kinase D (PKD), which phosphorylates CERT and OSBP to release these proteins from the Golgi complex^29,30^. PKD kinase activity also contributes to the membrane fission reaction required for transport carrier biogenesis at the TGN^31–33^.

On the basis of these reports and our previous data, we proposed a model in which lipid transfer at ER-Golgi contact sites promotes the biogenesis of carriers of the TGN to the cell surface (CARTS)^34^, which transport selective cargoes from the TGN to the PM^35,36^. In this context, lipid transfer at these contact sites needs to be strictly controlled on demand for TGN-to-PM transport, but the molecular mechanisms underlying this regulation remain largely elusive. In this study, we identified SCAP as a novel Sac1-interacting protein at ER-Golgi contact sites. Our data show a new function of SCAP under cholesterol-fed conditions, in which SCAP, together with SREBPs, interacts with the cholesterol/PI4P exchange machinery at ER-Golgi contact sites and facilitates the biogenesis of CARTS at the TGN in an ER cholesterol-dependent manner.

## Results

### Identification of SCAP as a novel component of Sac1-positive ER-Golgi contact sites

We previously reported that the perinuclear Sac1 localization represents its presence at specialized ER subdomains that are closely apposed to the TGN membranes, most likely corresponding to ER-Golgi contact sites^34^, rather than at the Golgi complex ^37^. To identify novel components of ER-Golgi contact sites, we explored Sac1-interacting proteins using a Sac1 K2A mutant, which is more highly enriched in the juxtanuclear region than the wild-type (WT) protein, as revealed by its reduced peripheral staining (Fig. 1a). This mutant has replacement of two lysines in the C-terminal COPI-interacting motif with alanines. While the di-lysine COPI-interacting motif is used for retrograde transport from the *cis*-Golgi to the ER^38^, some ER integral membrane proteins, including Bap31, also utilize this signal for their movement along the ER from the cell center to the periphery^39,40^. The deconvolved, enhanced contrast, and increased resolution images of cells treated with 25-hydroxycholesterol (25-HC), a reagent that binds to OSBP and promotes the targeting of the VAP/OSBP complex to ER-Golgi contact sites ^22,41,42^, showed good overlap between the green fluorescent protein (GFP)-Sac1 K2A signal and that of the integral ER membrane protein VAP-A, and also the close apposition, but not full overlap, of GFP-Sac1 K2A to the TGN marker TGN46, similarly to WT (Fig. 1b). Consistently, the continuity of GFP-Sac1 K2A-positive juxtanuclear membrane compartments with the peripheral ER membrane was demonstrated by inverse fluorescence recovery after photobleaching (iFRAP) (Fig. 1c): photobleaching of the peripheral area immediately decreased the perinuclear signals of GFP-Sac1 WT and K2A, but not of the Golgi resident protein N-acetylglucosaminyl transferase I-GFP (negative control). These results suggest that the Sac1 K2A mutant is prominently localized to ER-Golgi contact sites and validate the use of this mutant as a bait to identify interacting proteins at the contact sites.

**Fig. 1.**
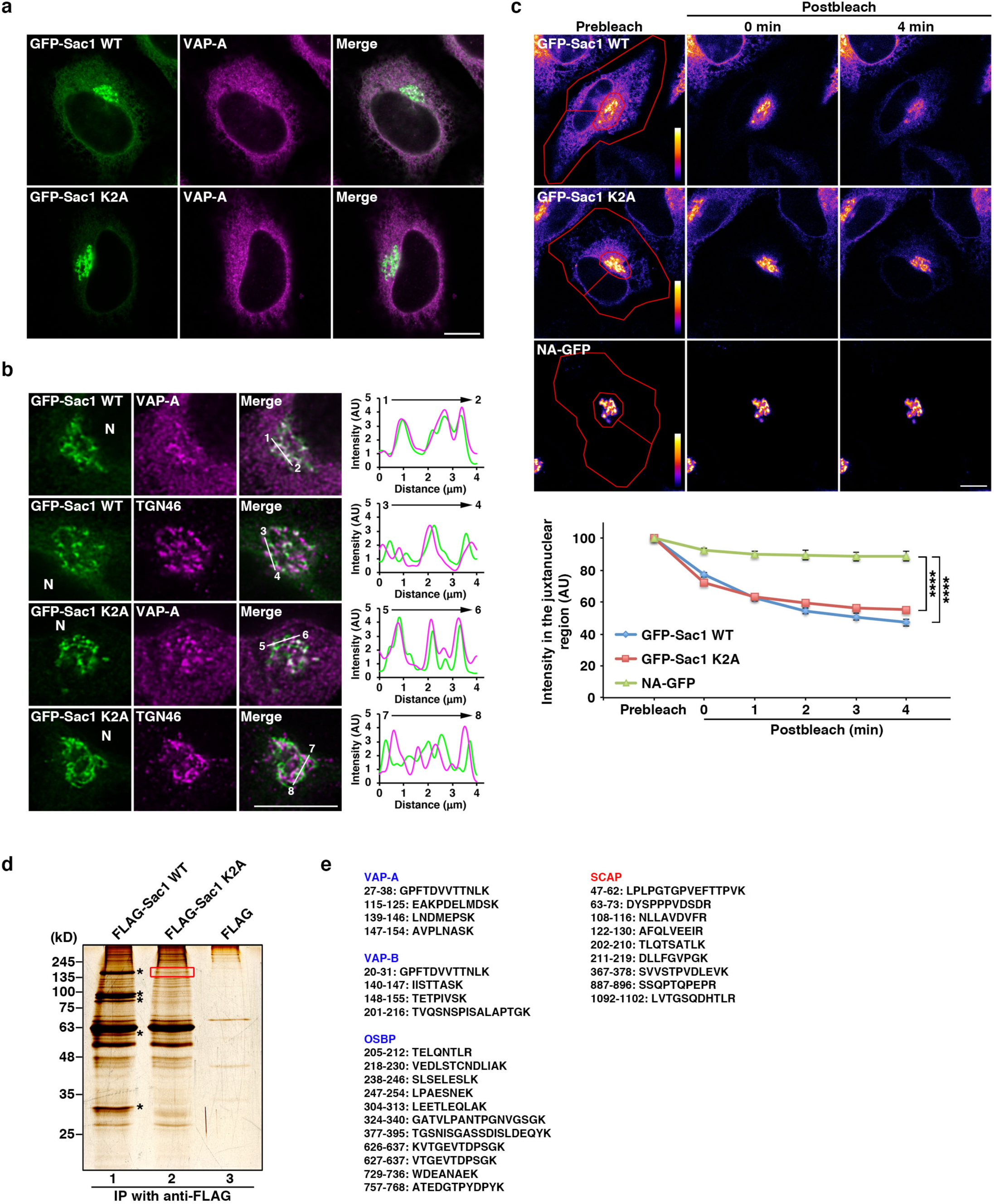
Identification of SCAP as a novel component of Sac1-positive ER-Golgi contact sites. **a**, Juxtanuclear localization of the GFP-Sac1 WT and K2A mutant in HeLa cells. **b**, Colocalization of GFP-Sac1 WT or K2A with VAP-A and their proximity localization with TGN46. HeLa cells expressing GFP-Sac1 WT or K2A were treated with 2 μg/mL 25-HC for 1 h. Images were subjected to deconvolution processing as described in *Materials and methods*. The graphs on the right show the fluorescence intensity of GFP-Sac1 WT or K2A (green), and VAP-A or TGN46 (magenta) along the respective white lines shown in the merged images. N, nucleus. **c**, iFRAP in HeLa cells expressing the GFP-Sac1 WT, K2A mutant, or N-acetylglucosaminyl transferase I (NA)-GFP. The areas delimited by a red line were bleached as described in *Materials and methods*. The graph below shows quantification of the fluorescence intensity of the indicated proteins in the non-bleached, juxtanuclear region. Data are means ± s.e.m. (n = 4 cells per condition, \*\*\*\**P* < 0.0001, one-way ANOVA multiple comparison test). **d**, Silver staining of immunoprecipitated proteins with the FLAG-Sac1 WT, K2A, or FLAG in HEK 293T cells. Asterisks denote protein bands containing COPI components. SCAP was identified in the protein band boxed with a red line. **e**, Peptides of VAP-A, VAP-B, OSBP, and SCAP, which were identified by mass spectrometric analysis of FLAG-Sac1 K2A immunoprecipitates (lane 2 in panel **d**). Scale bars, 10 μm.

Immunoprecipitation of FLAG-tagged Sac1 K2A followed by mass spectrometric analysis revealed multiple interactors of FLAG-Sac1 K2A (Fig. 1d, lane 2) with decreased binding of COPI coat proteins, which are major interactors for FLAG-Sac1 WT (Fig. 1d, lane 1, asterisks). The identified Sac1 K2A-interacting proteins include known components of ER-Golgi contact sites such as VAP-A, VAP-B, and OSBP, together with proteins with unknown functions at the contact sites, including SCAP (Fig. 1e). In here, we focus on SCAP because, although it had been previously shown to interact with VAP-A and VAP-B, the physiological role of this interaction remains unexplored^43^.

### ER-localized SCAP interacts through Sac1 with the VAP-A/OSBP complex at ER-Golgi contact sites

In order to confirm that SCAP is a bona fide component of ER-Golgi contact sites, the interactions of FLAG-tagged hamster SCAP with VAP-A, OSBP, and Sac1 were examined by immunoprecipitation (Fig. 2a). Endogenous Sac1 and VAP-A were clearly coprecipitated with FLAG-SCAP, but not with the FLAG vector control (Fig. 2a, lanes 7 and 9). When Myc-OSBP was coexpressed, this protein was coprecipitated with FLAG-SCAP and the interaction of FLAG-SCAP with VAP-A, but not with Sac1, was enhanced (Fig. 2a, lanes 9 and 10). As negative controls, we showed that an integral ER membrane protein, reticulon-4B (RTN-4B), and another lipid transfer protein, CERT, were not coprecipitated with FLAG-SCAP (Fig. 2b).

**Fig. 2.**
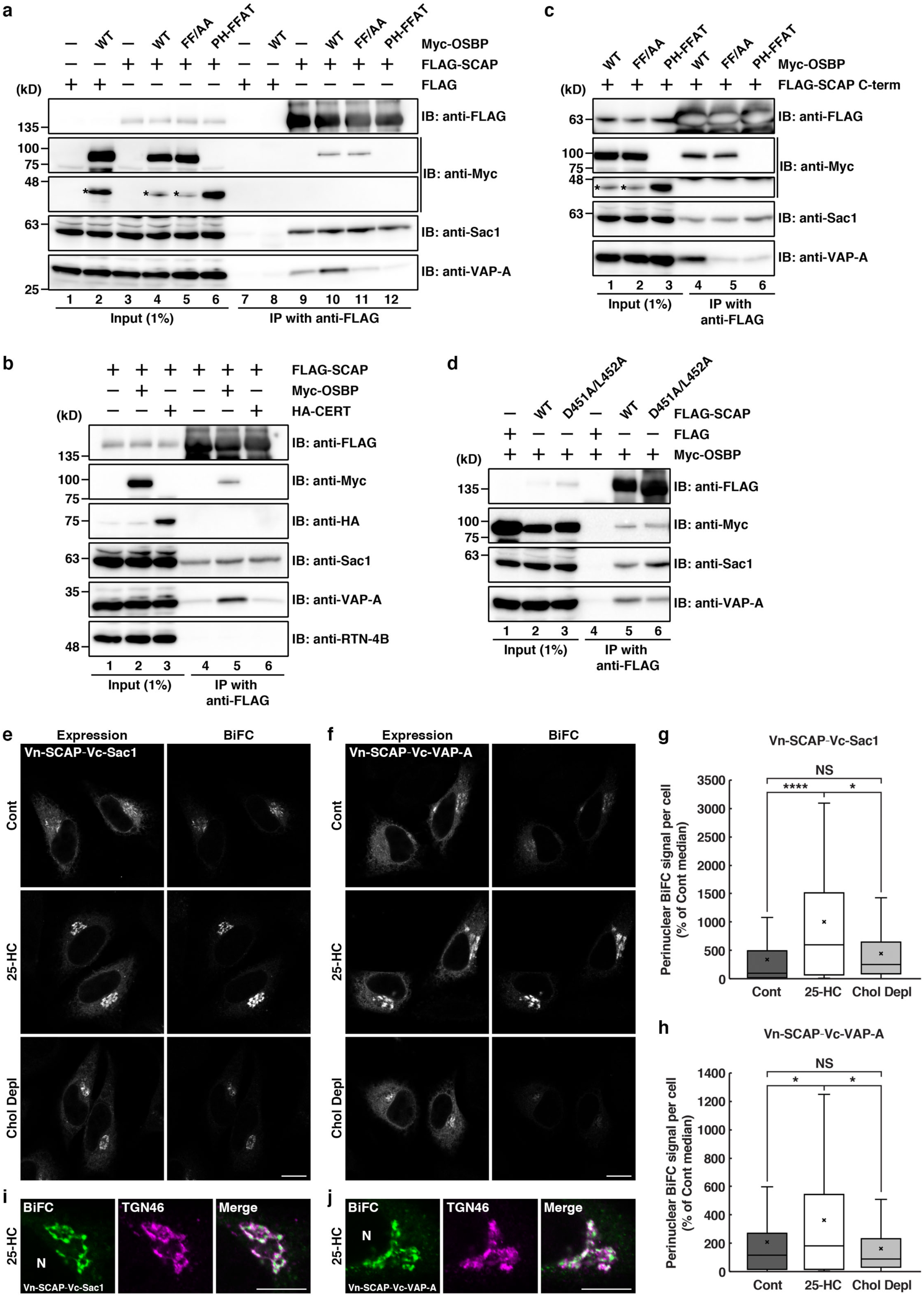
ER-localized SCAP interacts through Sac1 with the VAP-A/OSBP complex at ER-Golgi contact sites. **a**, Interactions of FLAG-SCAP with Myc-OSBP, Sac1, and VAP-A in HEK 293T cells. Cell lysates were subjected to immunoprecipitation with an anti-FLAG M2 affinity gel, and the cell lysates (Input) and immunoprecipitates (IP) were immunoblotted (IB) with the indicated antibodies. Asterisks denote degraded Myc-OSBP fragments. **b**, Interactions of FLAG-SCAP with Myc-OSBP, Sac1, and VAP-A, but not with HA-CERT or RTN-4B. **c**, Interactions of FLAG-SCAP C-term with Myc-OSBP, Sac1, and VAP-A. **d**, Interactions of the FLAG-SCAP D451A/L452A mutant with Myc-OSBP, Sac1, and VAP-A. **e**–**j**, BiFC visualization of SCAP/Sac1 or SCAP/VAP-A interactions at ER-Golgi contact sites in HeLa cells coexpressing Myc-OSBP and Venus N-terminal fragment (Vn)-fused SCAP in combination with Venus C-terminal fragment (Vc)-fused Sac1(**e**,**g**,**i**) or VAP-A (**f**,**h**,**j**). The cells were treated without [Control (Cont)] or with 2 μg/mL 25-HC for 2.5 h, or were cholesterol depleted (Chol Depl) for 3 h. The expression of Vn-SCAP together with Vc-Sac1 (**e**) or Vc-VAP-A (**f**) was visualized with an anti-GFP antibody. TGN46 was visualized with anti-TGN46 antibody (**i**,**j**). N, nucleus. The box-and-whisker plots show quantification of perinuclear BiFC signal (**g**,**h**). Boxes delimit the first and third quartiles and the central line is the median, whereas the cross represents the mean value. The whiskers represent the minimum and maximum values after outlier removal (Tukey whiskers) (Vn-SCAP/Vc-Sac1: control, n = 86 cells; 25-HC, n = 70 cells; cholesterol depleted, n = 101 cells, Vn-SCAP/Vc-VAP-A: control, n = 93 cells; 25-HC, n = 92 cells; cholesterol depleted, n = 101 cells, * P<0.05, \*\*\*\**P* < 0.0001, NS, not significant, Kruskal-Wallis multiple sample non-parametric test). Scale bars, 10 μm.

Next, we asked whether SCAP interacts with the VAP-A/OSBP complex via Sac1. Our previous work showed that Sac1 interacts with VAP-A via OSBP^34^. We used two OSBP mutants, FF/AA and PH-FFAT, that are defective in binding to VAP and Sac1, respectively (Ref^44,45^ and Supplementary Fig. 1a,b). In cells expressing Myc-OSBP FF/AA, this protein, but not VAP-A, was found to interact with FLAG-SCAP (Fig 2a, lane 11), suggesting that VAP-A is not needed for the interaction of SCAP with OSBP. By contrast, Myc-OSBP PH-FFAT was not coprecipitated with FLAG-SCAP (Fig 2a, lane 12). Although previous work revealed that Myc-OSBP PH-FFAT fixes and expands ER-Golgi contact sites through its stable interaction with VAP-A^22^, it inhibited the interaction of FLAG-Sac1 with VAP-A (Supplementary Fig. 1, lane 12). Myc-OSBP PH-FFAT defective in Sac1 binding also greatly reduced the interaction of FLAG-SCAP with VAP-A (Fig. 2a, lane 12), emphasizing the requirement of Sac1 for the interaction of SCAP with the VAP-A/OSBP complex. Similar results were obtained for a FLAG-tagged C-terminal cytoplasmic fragment (C-term) of SCAP^46^ (Fig. 2c) and the specific interaction between FLAG-SCAP and Sac1 was also detected in HeLa cells stably expressing FLAG-SCAP at a level of about four times the level of endogenous SCAP (Supplementary Fig. 1c,d). These results exclude the possibility of membrane aggregation or co-sedimentation with FLAG-SCAP. The specific but relatively weak interactions of SCAP with the cholesterol/PI4P exchange machinery most likely reflect transient interactions among these components, consistent with the underlying association-dissociation dynamics of ER-Golgi contacts that are crucial for their function^22,34^.

To corroborate that ER-localized SCAP, but not Golgi-localized SCAP, can form complexes with VAP-A, OSBP, and Sac1 at ER-Golgi contact sites, we used a SCAP mutant with replacement of aspartic acid 451 and leucine 452 with alanines at the MELADL motif in loop 6 (D451A/L452A), which is defective in COPII binding and therefore cannot exit the ER in COPII-coated vesicles^47^. FLAG-SCAP D451A/L452A showed interactions with VAP-A, OSBP, and Sac1, analogous to FLAG-SCAP WT (Fig. 2d). We next visualized the interactions of SCAP with these proteins in intact cells by using bimolecular fluorescence complementation (BiFC). As shown previously^48,49^, a BiFC signal derived from the Vn-OSBP/Vc-VAP-A interaction was detected at the perinuclear region representing ER-Golgi contact sites (Supplementary Fig. 2c, top row), whereas no signal was detected when only Vn- or Vc-fused constructs were individually expressed (Supplementary Fig. 2a,b). This signal was enhanced by 25-HC treatment (Supplementary Fig. 2c, middle row). Similar results were obtained with the combination of Vn-OSBP/Vc-Sac1 (Supplementary Fig. 2d). The Vn-SCAP/Vc-Sac1 and Vn-SCAP/Vc-VAP-A interactions were also observed upon coexpression of Myc-OSBP, and these interactions were enhanced by 25-HC treatment (Fig. 2e,f,g,h), similar to the Vn-OSBP/Vc-VAP-A and Vn-OSBP/Vc-Sac1 interactions (Supplementary Fig. 2c,d). Upon 25-HC treatment, the BiFC signal derived from the Vn-SCAP and Vc-Sac1 or Vc-VAP-A interaction was in close apposition to the TGN46 signal, confirming that these interactions occur at ER-Golgi contact sites (Fig. 2i,j).

To further substantiate that the perinuclear Vn-SCAP/Vc-Sac1 and Vn-SCAP/Vc-VAP-A BiFC signals represent the interaction of ER-localized SCAP, but not of Golgi-localized SCAP, we examined the effect of cholesterol depletion. As reported previously^50^, cholesterol depletion using lipoprotein-deficient serum and 2-hydroxypropyl-β-cyclodextrin caused redistribution of a part of the SCAP pool from the ER to the *cis/medial* Golgi membranes (Supplementary Fig. 3a,b) accompanied by SREBP2 cleavage (Supplementary Fig. 3c). In contrast to 25-HC treatment, cholesterol depletion did not enhance either the Vn-SCAP/Vc-Sac1 or Vn-SCAP/Vc-VAP-A interaction (Fig. 2e,f, bottom row, and Fig 2g,h). Altogether, these results suggest that ER-localized SCAP preferentially interacts with VAP-A, OSBP, and Sac1 at ER-Golgi contact sites.

### SCAP is important for PI4P turnover and VAP-A/OSBP complex distribution at ER-Golgi contact sites

The finding that SCAP forms complexes with VAP-A, OSBP, and Sac1 at ER-Golgi contact sites prompted us to examine whether SCAP is involved in the counter-transport of ER cholesterol and Golgi PI4P at ER-Golgi contact sites. To address this issue, we performed knockdown of SCAP in HeLa cells. By using siRNA, the expression of SCAP was decreased to ∼20% of the control level (Fig. 3a). In response to cholesterol deprivation, the SCAP-SREBP pathway stimulates transcription of genes responsible for cholesterol synthesis and uptake, such as 3-hydroxy-3-methylglutaryl coenzyme A reductase (HMGR) and low-density lipoprotein receptor (LDLR)^16,51^. However, under cholesterol-fed conditions –that is, culturing of cells in normal medium supplemented with fetal calf serum (FCS) as a source of sterols–, SCAP knockdown did not reduce the transcription of either HMGR or LDLR genes, but rather slightly increased the LDLR mRNA levels (Fig. 3b). Consistently, no significant change in the total cholesterol level was observed upon SCAP knockdown (Fig. 3c). Similar results were obtained for a HeLa stable cell line (shSCAP cells) with reduced expression of SCAP (∼10% of that in parental HeLa cells), but not of HMGR and LDLR (Fig. 3d,e,f).

**Fig. 3.**
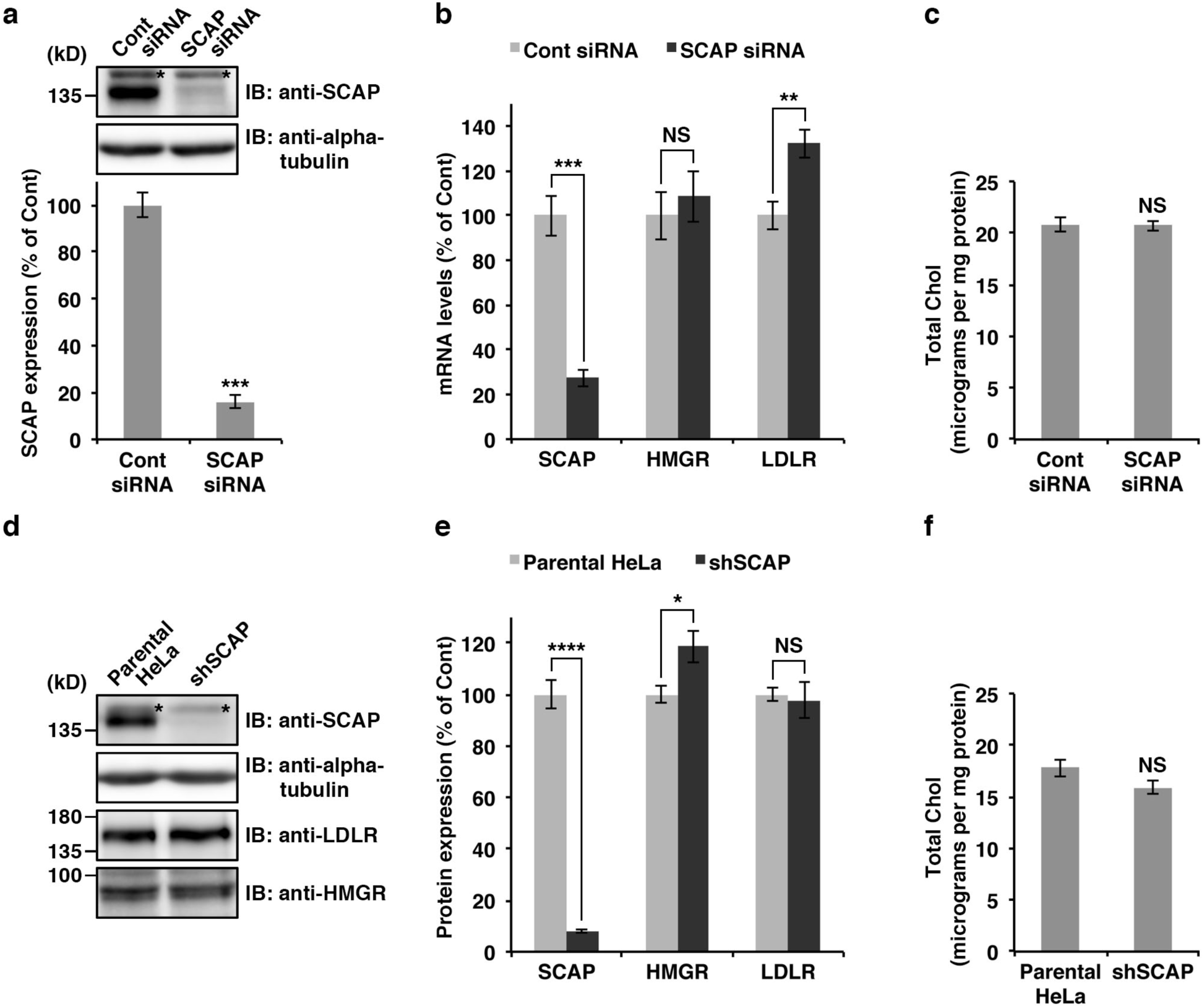
SCAP knockdown does not disrupt cholesterol metabolism under cholesterol-fed conditions. **a**, siRNA-mediated knockdown of SCAP in HeLa cells, monitored by Western blotting. The graph shows determination of the expression levels of SCAP at 72 h after siRNA transfection. Data are means ± s.e.m. (n = 3 independent experiments, \*\*\**P* < 0.001, unpaired two-tailed Student’s *t*-test). Asterisks on the Western blot denote nonspecific bands. **b**, Determination of the mRNA levels of the indicated genes in control (Cont) and SCAP knockdown cells by quantitative real-time PCR. Data are means ± s.e.m. (n = 4 independent experiments, \*\**P* < 0.01, \*\*\**P* < 0.005, NS, not significant, unpaired two-tailed Student’s *t*-test). **c**, Total cholesterol (Chol) levels in control and SCAP knockdown cells. Data are means ± s.e.m. (n = 6 independent experiments, NS, not significant, unpaired two-tailed Student’s *t*-test). **d**,**e**, shRNA-mediated knockdown of SCAP in HeLa cells, monitored by Western blotting. The graph (**e**) shows determination of the expression levels of SCAP, HMGR, and LDLR in parental HeLa (control) and shSCAP HeLa cells, as measured by Western blotting. Data are mean ± s.e.m. (n = 4 independent experiments, \**P* < 0.05, \*\*\*\**P* < 0.001, NS, not significant, unpaired two-tailed Student’s *t*-test). Asterisks on the Western blot denote nonspecific bands. **f**, Total cholesterol levels in parental HeLa and shSCAP HeLa cells. Data are means ± s.e.m. (n = 3 independent experiments, NS, not significant, unpaired two-tailed Student’s *t*-test).

Staining of parental HeLa and shSCAP cells with the fluorescent cholesterol probe filipin showed no significant difference in the intracellular distribution of cholesterol between these cell lines (Fig. 4a). Although a part of the perinuclear signal of filipin overlapped with that of a TGN marker, Golgin-97, most of the signal was thought to be derived from other membranous structures, as shown previously ^21^, and filipin does not seem to have sufficient sensitivity to detect low levels of cholesterol in ER and Golgi membranes.

**Fig. 4.**
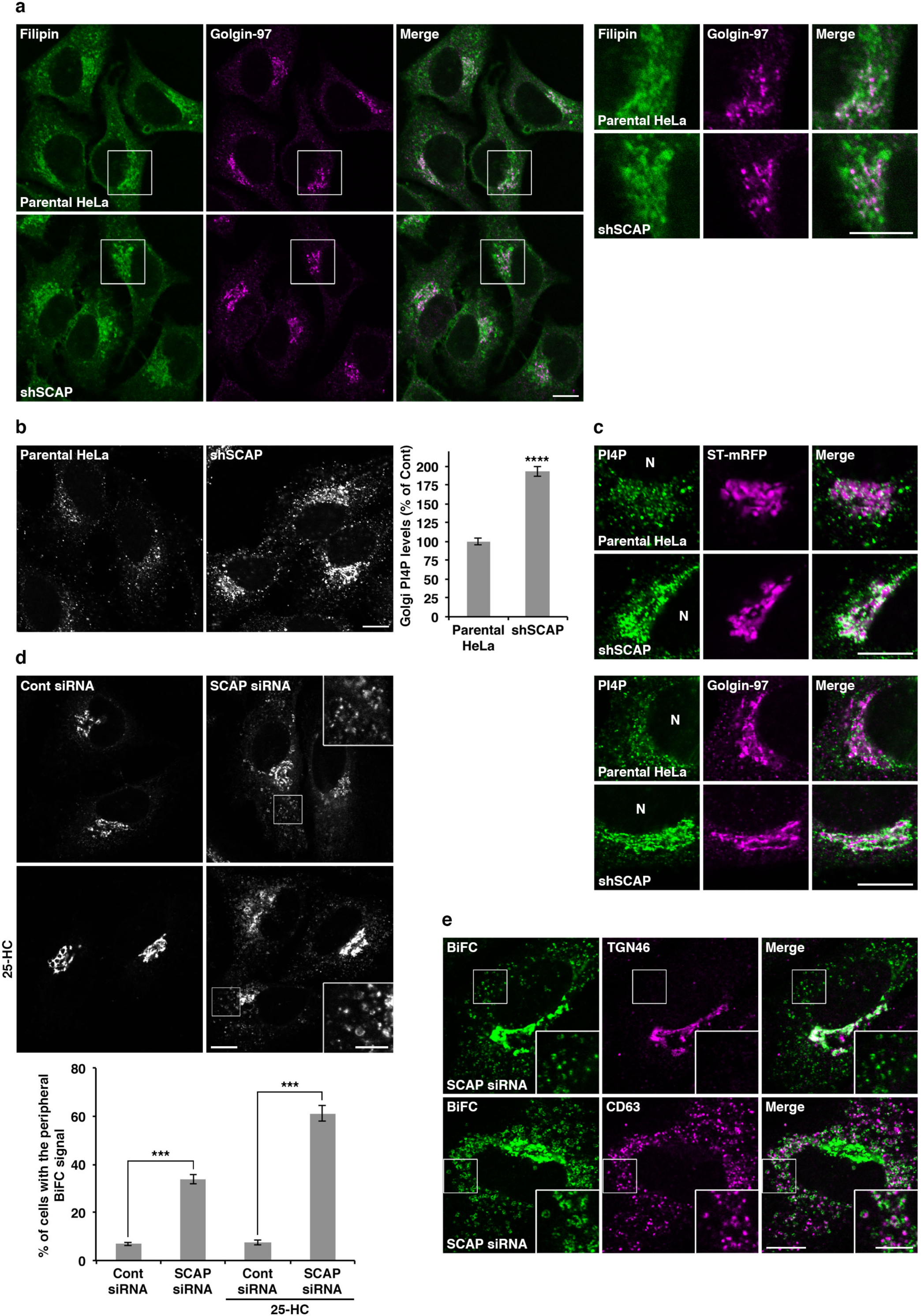
SCAP is important for PI4P turnover and VAP-A/OSBP complex distribution at ER-Golgi contact sites. **a**, Filipin staining in parental HeLa and shSCAP HeLa cells. High magnifications of the boxed areas are shown in the right panels. Scale bars, 10 μm. **b**,**c**, PI4P staining in parental HeLa and shSCAP HeLa cells with (top two rows in **c**) or without (**b** and bottom two rows in **c**) sialyltransferase (ST)-mRFP expression. N, nucleus. Scale bars, 10 μm. The graph shows determination of the Golgi PI4P levels in parental HeLa (control) and shSCAP HeLa cells. Data are means ± s.e.m. (n = 139 cells per conditions, \*\*\*\**P* < 0.0001, unpaired two-tailed Student’s *t*-test). **d**, BiFC visualization of OSBP/VAP-A interactions in control (Cont) and SCAP knockdown cells. HeLa cells stably coexpressing Vn-OSBP and Vc-VAP-A were transfected with siRNA. After 72 h, the cells were treated with or without 2 μg/mL 25-HC for 2.5 h. High magnifications of the boxed areas are shown in the insets where brightness/contrast-enhancement was applied. Scale bars, 10 μm (large panels), 5 μm (insets). The graph shows the percentage of cells with the peripheral BiFC signal of Vn-OSBP/Vc-VAP-A. Data are means ± s.e.m. (n = 3 independent experiments, 100-131 cells per condition, \*\*\**P* < 0.005, unpaired two-tailed Student’s *t*-test). **e**, Close apposition of CD63-but not of TGN46-positive membranes to the peripheral BiFC signal of Vn-OSBP/Vc-VAP-A in SCAP knockdown cells treated with 2 μg/mL 25-HC for 2.5 h. High magnifications of the boxed areas are shown in the insets. Scale bars, 10 μm (large panels), 5 μm (insets).

Because of the lack of cholesterol probes with enough sensitivity for our purposes, we focused on monitoring PI4P level in SCAP knockdown cells. When PI4P was visualized with a specific antibody, the fluorescence signal in shSCAP cells was significantly increased as compared to that in the parental HeLa cells, especially in the juxtanuclear region that partially overlapped with membranes positive for the TGN markers sialyltransferase and Golgin-97 (Fig. 4b,c). Similar phenotypes were observed upon VAP-A/B or CERT/OSBP double knockdown, as well as in Sac1 knockdown cells (Supplementary Fig. 4a). Taken together, our data suggest that SCAP regulates the TGN PI4P levels by controlling the turnover of PI4P at ER-Golgi contact sites.

Next, we evaluated the effect of SCAP knockdown on the formation of the VAP-A/OSBP complex by using the BiFC approach. A stable cell line coexpressing Vn-OSBP and Vc-VAP-A showed a BiFC signal in the juxtanuclear region, which was enhanced by 25-HC treatment (Supplementary Fig. 4b). When SCAP was knocked down by siRNA, ∼34% of cells showed a bright BiFC signal in the peripheral region, in addition to the signal at the juxtanuclear Golgi region, and the number of cells with this phenotype increased to ∼62% upon 25-HC treatment (Fig. 4d). As the VAP/OSBP complex has been reported to exist not only at ER-Golgi contact sites, but also at ER-endosome contact sites^52^, we compared the localization of the peripheral BiFC signal with that of the late endosomal marker CD63. The close apposition of CD63 but not of TGN46 to the peripheral BiFC signal was observed in SCAP knockdown cells (Fig. 4e), indicating that SCAP is required to prevent the redistribution of the VAP-A/OSBP complex to ER-endosome contact sites and to maintain it at ER-Golgi contact sites.

### SCAP is required for the biogenesis of CARTS at the TGN

We previously reported that perturbation of lipid transfer complexes at ER-Golgi contact sites inhibits the biogenesis of CARTS at the TGN^34^. We now visualized CARTS in Vn-OSBP/Vc-VAP-A expressing cells treated with 25-HC by staining a CARTS specific cargo, pancreatic adenocarcinoma up-regulated factor (PAUF)^35^. Deconvolved images showed that putative nascent CARTS were located in the close vicinity of BiFC-positive perinuclear region (Fig. 5a), suggesting that CARTS form at sites immediately adjacent to the VAP-A/OSBP-containing ER-Golgi contact sites. Next, we investigated whether SCAP is required for CARTS-mediated protein secretion. In SCAP knockdown cells, the secretion of PAUF-MycHis was reduced to ∼50% of that in control cells (Fig. 5b). Moreover, the small amount of PAUF-MycHis secreted by SCAP knockdown cells was detected as a smeared band, suggesting a defect in PAUF processing, as previously observed in cells depleted of other ER-Golgi contact site components^34^.

**Fig. 5.**
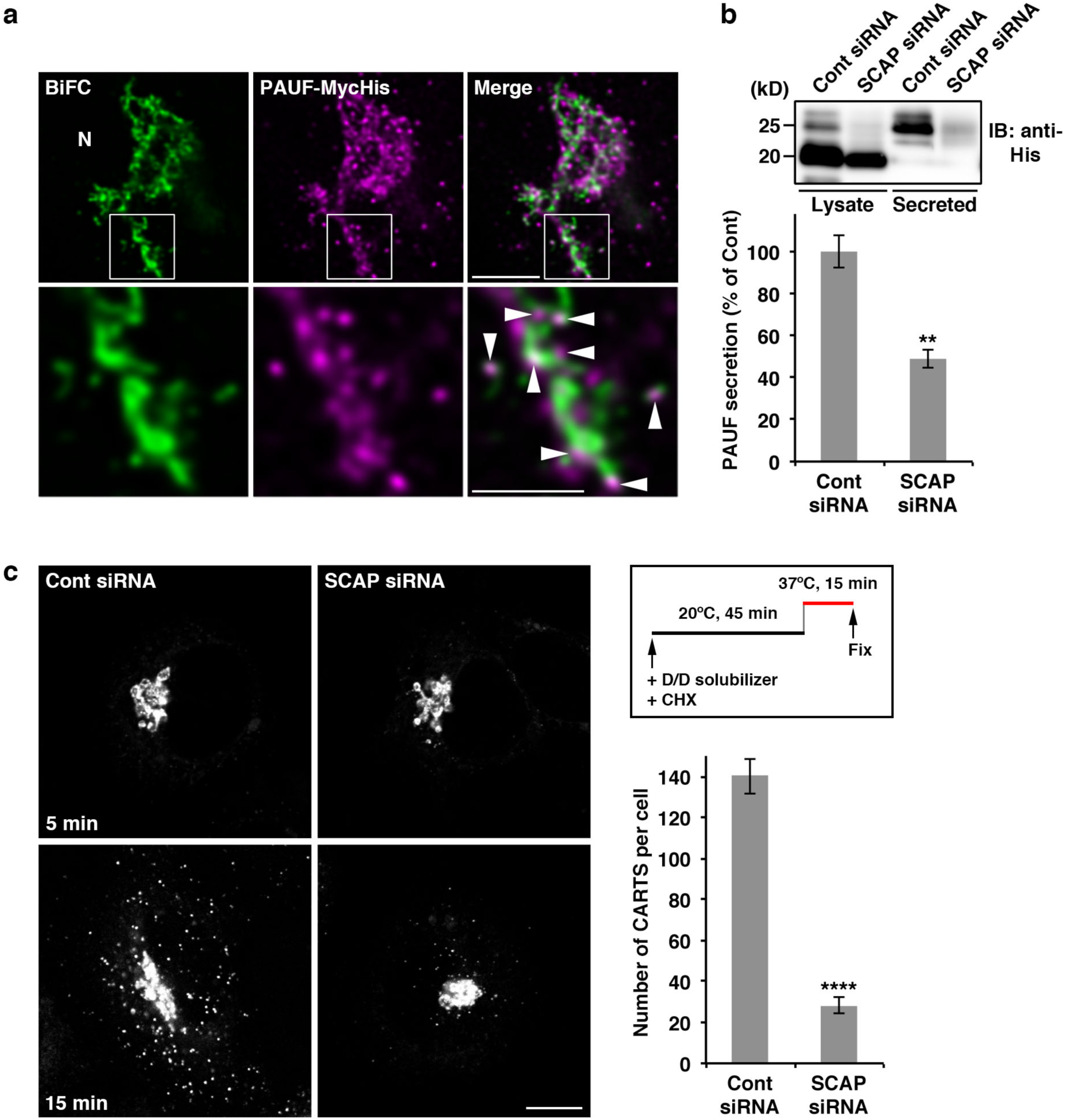
SCAP is required for the biogenesis of CARTS at the TGN. **a**, Close proximity of CARTS formation sites to VAP-A/OSBP-mediated ER-Golgi contact sites. HeLa cells stably coexpressing Vn-OSBP and Vc-VAP-A were transfected with a plasmid for PAUF-MycHis. After 20 h, the cells were treated with 2 μg/mL 25-HC for 2.5 h. Images were subjected to deconvolution processing as described in *Materials and methods*. N, nucleus. High magnifications of the boxed areas are shown in the bottom row. Arrowheads indicate putative nascent CARTS located in the close vicinity of the BiFC signal of Vn-OSBP/Vc-VAP-A. Scale bars, 5 μm (upper row), 2.5 μm (bottom row). **b**, PAUF-MycHis secretion in control (Cont) and SCAP knockdown cells, monitored by Western blotting. The graph shows quantification of secreted PAUF-MycHis relative to the total cellular level and normalized as the values in control cells. Data are means ± s.e.m. (n = 3 independent experiments, \*\**P* < 0.01, unpaired two-tailed Student’s *t*-test). **c**, Biogenesis of mKate2-FM4-PAUF-containing CARTS in control and SCAP knockdown cells. The cells were incubated at 20°C with the D/D solubilizer and cycloheximide (CHX) for 45 min, followed by incubation at 37°C for 5 or 15 min. The graph shows the number of mKate2-FM4-PAUF-containing CARTS in control and SCAP knockdown cells at 15 min after the temperature shift to 37°C. Data are means ± s.e.m. (control siRNA: n = 20 cells; SCAP siRNA: n = 29 cells, \*\*\*\**P* < 0.0001, unpaired two-tailed Student’s *t*-test). Scale bar, 10 μm.

The effect of SCAP knockdown on the biogenesis of CARTS was assessed with an inducible CARTS formation assay, where synchronized transport of PAUF from the ER is carried out by using a reverse dimerization system involving D/D solubilizer-induced disassembly of FM4 domains^53^. The chimera protein mKate2-FM4-PAUF initially retained in the ER was first exported from this organelle and accumulated in the Golgi membranes by D/D solubilizer treatment at 20°C for 45 min, after which the temperature was shifted to 37°C to induce the formation of mKate2-FM4-PAUF-containing CARTS at the TGN. Live cell imaging showed that a large number of CARTS were formed at the TGN membranes and dispersed throughout the cytoplasm (Supplementary Video 1). Next, mKate2-FM4-PAUF was expressed in control and SCAP knockdown cells. At 15 min after the temperature shift to 37°C, the average number of CARTS in SCAP knockdown cells was ∼20% of that in control cells (Fig. 5c).

In our model, lipid transfer at ER-Golgi contact sites is thought to promote CARTS biogenesis through organization of cholesterol- and SM-enriched nanodomains at the TGN^34^. We therefore examined the effect of SCAP knockdown on transport of glycosylphosphatidyl inositol (GPI)-anchored protein, which has been reported to associate with such lipid nanodomains^1,54,11,55^. Synchronized transport of mKate2-FM4-GPI from the ER to the PM was initiated by addition of the D/D solubilizer, as previously reported^11^. SCAP knockdown did not affect transport of mKate2-FM4-GPI from the ER (0 min) to the Golgi complex (30 min). Nevertheless, at 60 min after transport initiation, the amount of mKate2-FM4-GPI that had reached the cell surface was significantly decreased in SCAP knockdown cells (Fig. 6a). At 90 min most of the protein had been transported to the PM in control cells, but in ∼40% of SCAP knockdown cells, the protein was still localized to the TGN (Fig. 6a,b). Similar results were obtained with knockdown of VAP-A/B, CERT/OSBP, or, to a lesser extent, OSBP (Fig. 6a and Supplementary Fig. 5a). In addition to these results, overexpression of OSBP PH-FFAT, which immobilizes ER-Golgi contact sites and inhibits CARTS biogenesis^34^, strongly inhibited mKate2-FM4-GPI export from the TGN (Supplementary Fig. 5b).

**Fig. 6.**
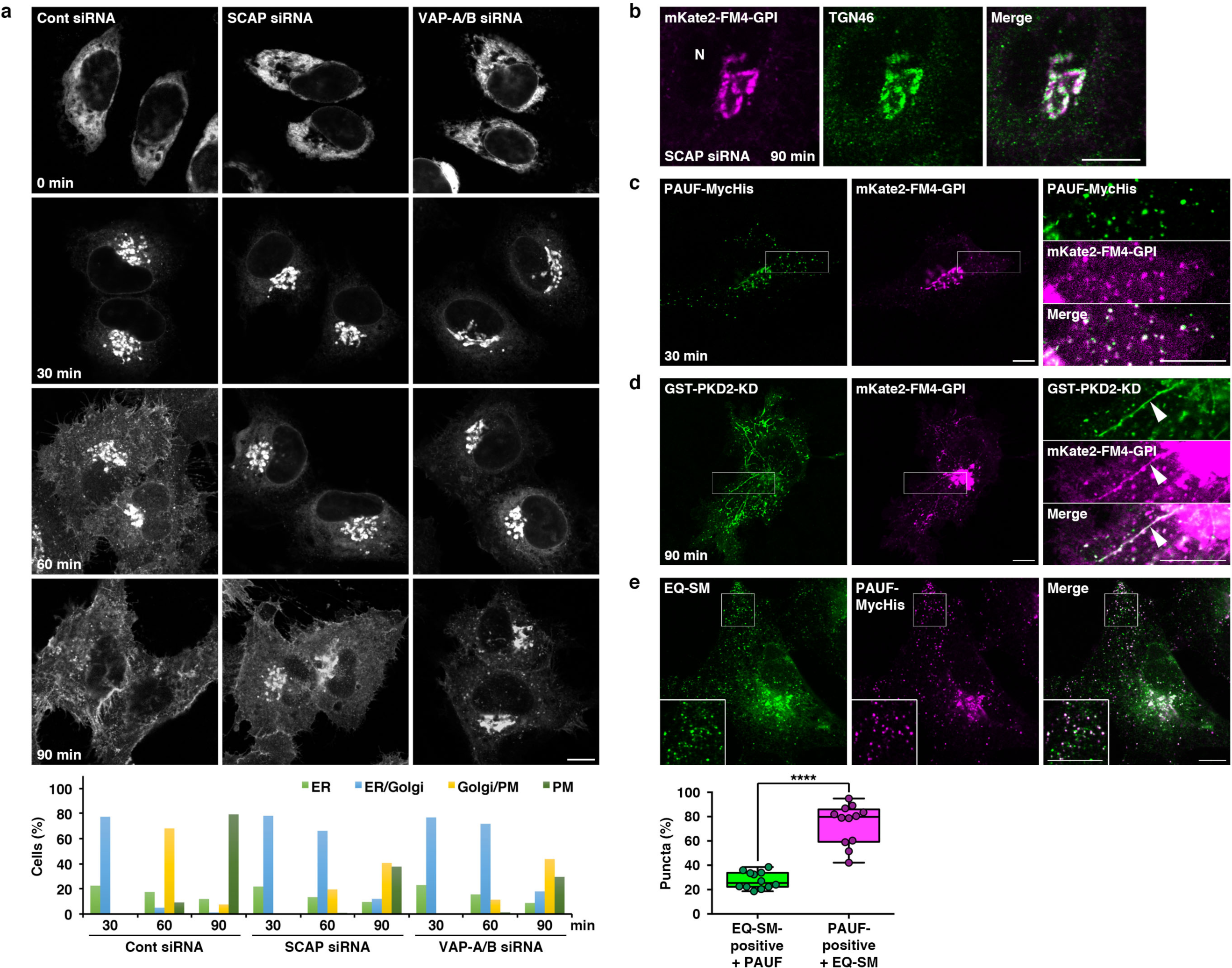
SCAP is required for GPI-anchored protein transport from the TGN to the PM. **a**, mKate2-FM4-GPI transport from the ER to the PM via the Golgi complex in control, SCAP, and VAP-A/B knockdown cells. The cells were incubated at 37°C with the D/D solubilizer and cycloheximide, and fixed at the indicated times. The graph shows the percentages of cells with mKate2-FM4-GPI at the ER, ER/Golgi, Golgi/ PM, or PM at the indicated times. The data shown are for a single representative experiment out of three performed (control siRNA: 30 min, n = 235 cells; 60 min, n = 239 cells; 90 min, n = 252 cells; SCAP siRNA: 30 min, n = 248 cells; 60 min, n = 246 cells; 90 min, n = 241 cells; VAP-A/B siRNA: 30 min, n = 230 cells; 60 min, n = 238 cells; 90 min, n = 240 cells). **b**, mKate2-FM4-GPI has accumulated at the TGN at 90 min after the transport induction in SCAP knockdown cells. N, nucleus. **c**, Colocalization of PAUF-MycHis and mKate2-FM4-GPI in CARTS at 30 min after transport induction. **d**, Colocalization of GST-PKD2-kinase dead (KD) and mKate2-FM4-GPI in tubules attached to the TGN at 90 min after transport induction. In **c** and **d**, high magnifications of the boxed areas are shown in the right column where brightness/contrast-enhancement was applied to the mKate2-FM4-GPI channel. Arrowheads in **d** indicate a GST-PKD2-KD-induced tubule containing mKate2-FM4-GPI. **e**, Colocalization of EQ-SM (tagged with oxGFP) and PAUF-MycHis. High magnifications of the boxed areas are shown in the insets. The box-and-whisker plots show quantification of EQ-SM-positive puncta containing PAUF-MycHis (green) and PAUF-MycHis-positive puncta (CARTS) containing EQ-SM (magenta). Boxes delimit the first and third quartiles and the central line is the median. The whiskers represent the minimum and maximum values after outlier removal (Tukey whiskers) (EQ-SM-positive: n = 4487 puncta, PAUF-positive: n=1650 puncta in 12 cells, \*\*\*\**P* <0.0001, paired two-tailed Student’s *t*-test). Scale bars, 10 μm.

We tested if CARTS are the carriers that transport mKate2-FM4-GPI from the TGN to the cell surface. When mKate2-FM4-GPI and PAUF-MycHis were coexpressed in cells, we observed that mKate2-FM4-GPI was included in PAUF-MycHis-positive CARTS, although its fluorescence signal was relatively low (Fig. 6c). Consistent with our previous finding that CARTS biogenesis requires PKD-mediated membrane fission at the TGN^35^, export of mKate2-FM4-GPI from the TGN was inhibited by expression of a dominant-negative kinase-dead mutant of PKD2 and even at 90 min after transport initiation, the protein was included in TGN-derived tubules, most likely corresponding to fission-defective precursors of transport carriers, such as CARTS (Fig. 6d). Previous work revealed that a non-toxic SM reporter protein, EQ-SM is enriched in transport carries containing mKate2-FM4-GPI^11^. When EQ-SM and PAUF-MycHis were coexpressed in cells, ∼80% of PAUF-MycHis-positive CARTS contained EQ-SM (Fig. 6e). This result fits well with the former report that 86 ± 5% of mKate2-FM4-GPI-positive exocytic vesicles contain EQ-SM^11^. Taken together, these results strongly suggest that SCAP promotes the biogenesis of CARTS, which are enriched in cholesterol and SM.

### SCAP regulates CARTS biogenesis in a cholesterol-dependent manner

SCAP directly binds cholesterol and plays a pivotal role in cholesterol homeostasis as a cholesterol sensor in the ER membrane^20,56,57^. Does SCAP regulate CARTS biogenesis by sensing the ER cholesterol level? Previous mutagenesis analysis of SCAP in Chinese hamster ovary (CHO) cells revealed that several amino acid residues are critical for the conformational change induced by cholesterol binding^20^. Specifically, replacement of tyrosine 234 in the cholesterol-binding loop1 region of SCAP (Fig. 7a) with alanine (Y234A) abolishes binding of loop1 to loop7, and thus this mutant binds to Insig even in the absence of cholesterol^57^. By contrast, a mutant with replacement of tyrosine 298 in the transmembrane sterol-sensing domain (Fig. 7a) with cysteine (Y298C) is resistant to the cholesterol-induced conformational change and shows no Insig binding even in the presence of sterols^43,58–61^. We established different shSCAP cells, each of which stably expressing hamster SCAP WT, Y234A, or Y298C. Western blotting of cell lysates with an antibody specific to hamster SCAP, and one recognizing both human and hamster SCAP indicated that Y234A was less expressed than WT and Y298C, but it was still expressed at a considerably higher level than the endogenous protein in parental HeLa cells (Fig. 7b). Immunofluorescent staining with an anti-hamster SCAP antibody showed reticular distributions of WT, Y234A, and Y298C in the ER, under cholesterol-fed conditions (Supplementary Fig. 3d, upper row). In a previous work, digestion of *N*-linked carbohydrates of SCAP by endoglycosidase (endo) H was shown to indicate that SCAP Y298C is redistributed to the Golgi complex even in the presence of sterols^58^. However, at the level of immunofluorescence microscopy, the signal of Y298C in the Golgi complex was not obvious, because most of the protein was present in the juxtanuclear ER (Supplementary Fig. 3d, upper right panel). Perhaps, a minor fraction of the Y298C pool is transported to the Golgi complex and acquires endo H resistance in the presence of sterols. When cholesterol was depleted from cells, fractions of the WT and Y298C pools were redistributed from the ER to the Golgi complex, but Y234A remained in the ER (Supplementary Fig. 3d, lower row), in agreement with previous reports^57,58,60^.

**Fig. 7.**
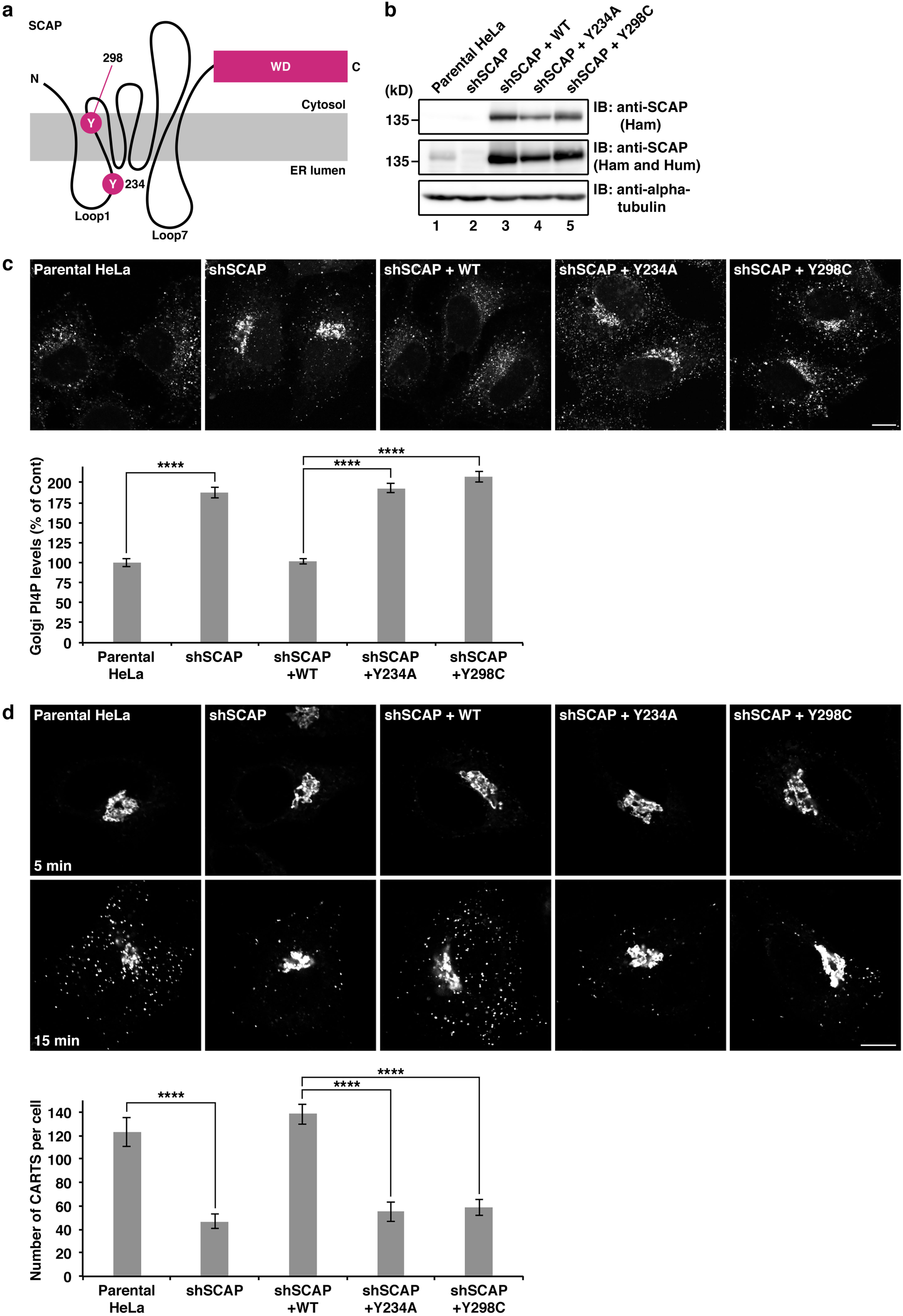
SCAP regulates CARTS biogenesis in a cholesterol-dependent manner. **a**, Schematic representation of the SCAP topology, where the Y234 and Y298 residues, loop1 and loop7, and WD (tryptophan–aspartate) repeats are indicated. **b**, Establishment of shSCAP HeLa cells stably expressing the hamster SCAP WT, Y234A, or Y298C mutants. Cell lysates were immunoblotted (IB) with the indicated antibodies. Ham, hamster; Hum, human. **c**, Recovery of PI4P turnover on expression of hamster SCAP WT, but not of Y234A or Y298C. The graph shows determination of the Golgi PI4P levels in the indicated cells. Data are mean ± s.e.m. (Parental HeLa: n = 69 cells; shSCAP: n = 83 cells; shSCAP + WT: n = 78 cells; shSCAP + Y234A: n = 75 cells; shSCAP + Y298C: n = 70 cells, \*\*\*\**P* < 0.0001, one-way ANOVA multiple comparison test). **d**, Recovery of biogenesis of mKate2-FM4-PAUF-containing CARTS on expression of hamster SCAP WT, but not of Y234A or Y298C. The graph shows the number of mKate2-FM4-PAUF-containing CARTS in the indicated cells at 15 min after the temperature shift to 37°C. Data are means ± s.e.m. (n = 20 cells per conditions, \*\*\*\**P* < 0.0001, one-way ANOVA multiple comparison test). Scale bars, 10 μm.

By using these cell lines, we first examined the effect that the expression of the respective hamster SCAP proteins has on the turnover of PI4P. Our results showed that expression of WT, but not Y234A or Y298C, reversed the accumulation of PI4P at the TGN (Fig. 7c). We next examined their effects on CARTS biogenesis. Expression of SCAP WT caused the recovery of the number of CARTS from SCAP knockdown, but neither Y234A nor Y298C showed such an effect (Fig. 7d). These results suggest that the sterol-sensing ability of SCAP is required for CARTS biogenesis, tightly linked to the capacity of SCAP to regulate PI4P turnover at ER-Golgi contact sites.

### SCAP/SREBP complex functions in CARTS biogenesis

SCAP and SREBPs are known to form a stable complex. Consistent with previous reports^62–64^, expression levels of SREBP1 and SREBP2 were also significantly decreased in shSCAP cells (Fig. 8a, lane 2). Importantly, while expression of not only SCAP WT, but also Y234A or Y298C recovered expression of SREBPs (Fig. 8a, lanes 3-5), these mutants were not able to rescue the defects in PI4P turnover and CARTS biogenesis (Fig. 7c,d). These results suggest that the sterol-sensing ability of SCAP, and not the presence of SREBPs, is the dominant factor by which SCAP controls PI4P turnover and CARTS biogenesis. Immunoprecipitation of FLAG-SREBP1a and FLAG-SREBP2 showed their interactions with Myc-OSBP and endogenous Sac1 and VAP-A (Fig. 8b). Our observation that FLAG-SREBPs show a higher affinity for GFP-SCAP than for components of the cholesterol/PI4P exchange machinery (Fig. 8c), suggests that SCAP/SREBPs form a stable complex that only transiently interacts with the lipid exchange machinery at ER-Golgi contact sites. Finally, we tested whether knockdown of SREBPs affects CARTS biogenesis. The expression levels of SREBP1 and SREBP2 was significantly decreased by using siRNA targeting common sequence of two alternative splicing isoforms of SREBP1 (1a and 1c) and SREBP2, respectively, without affecting expression of SCAP (Fig. 8d, lanes 1-3). Since a mixture of these two siRNAs did not reduce SREBP1 and SREBP2 at the same time (Fig. 8d, lane 4), the effect on CARTS formation of either SREBP1 or SREBP2 knockdown was examined. Our results showed that the average number of CARTS was decreased to ∼71% and ∼36% of the control levels in SREBP1 and SREBP2 knockdown cells, respectively (Fig. 8e).

**Fig. 8.**
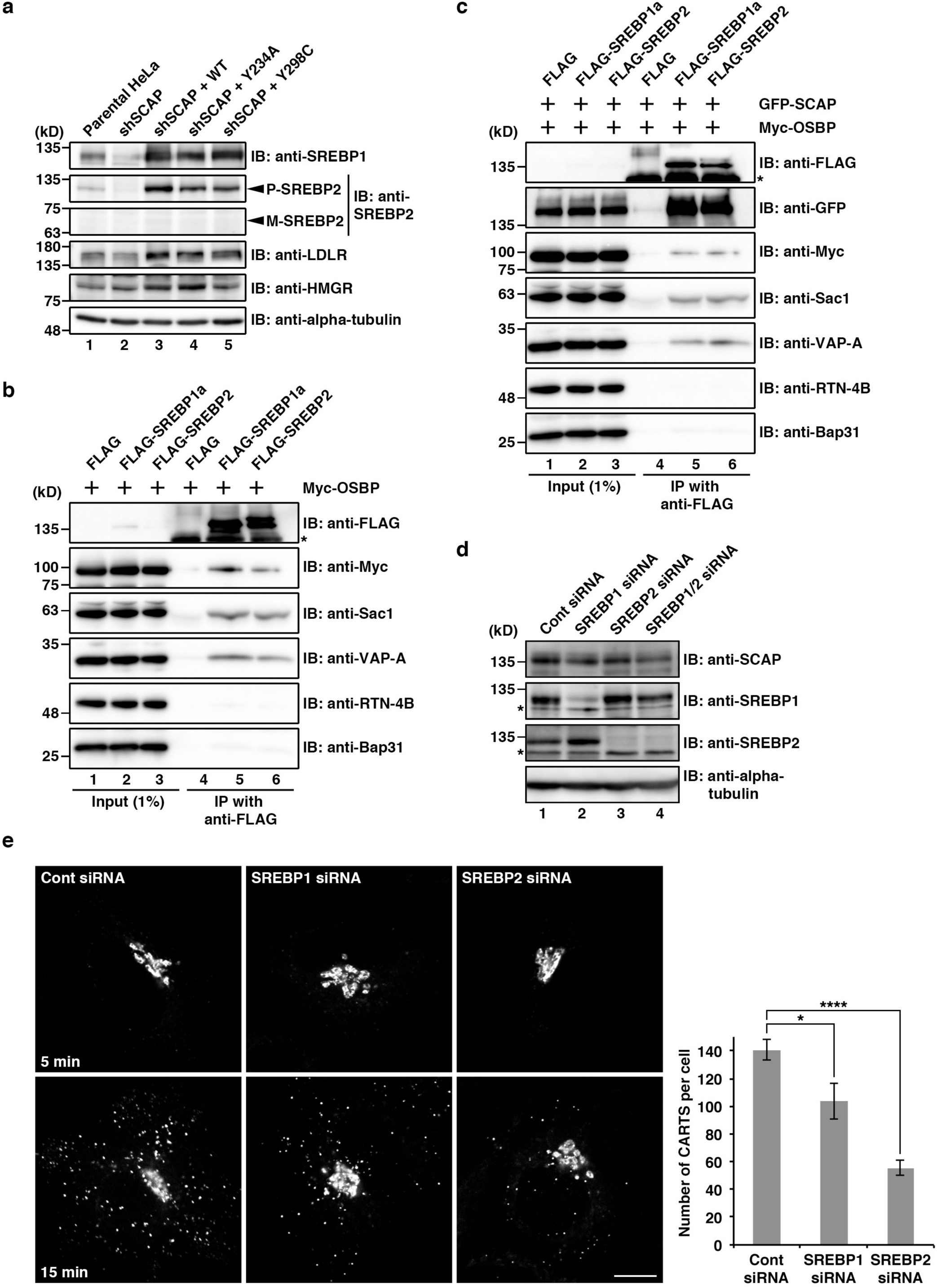
SCAP/SREBP complex functions in CARTS biogenesis. **a**, Effects of SCAP knockdown and expression of the hamster SCAP WT, Y234A, or Y298C mutants on the expression levels of SREBP1 and SREBP2. Cell lysates were immunoblotted (IB) with the indicated antibodies. The precursor (P), but not mature (M) forms of SREBP2 were detected in parental HeLa cells and shSCAP HeLa cells stably expressing the hamster SCAP WT, Y234A, or Y298C mutants. Expression levels of LDLR and HMGR were slightly increased by expression of hamster SCAP WT, Y234A, or Y298C mutants. **b**, Interactions of FLAG-SREBP1a and FLAG-SREBP2 with Myc-OSBP, Sac1, and VAP-A, but not with RTN-4B and Bap31 in HEK 293T cells. Cell lysates were subjected to immunoprecipitation with an anti-FLAG M2 affinity gel, and the cell lysates (Input) and immunoprecipitates (IP) were immunoblotted (IB) with the indicated antibodies. Asterisk denotes nonspecific bands. **c**, Interactions of FLAG-SREBP1a and FLAG-SREBP2 with GFP-SCAP, Myc-OSBP, Sac1, and VAP-A, but not with RTN-4B and Bap31. Asterisk denotes nonspecific bands. **d**, siRNA-mediated knockdown of SREBP1 and/or SREBP2 in HeLa cells, monitored by Western blotting. Asterisks denote nonspecific bands. **e**, Biogenesis of mKate2-FM4-PAUF-containing CARTS in control, SREBP1, and SREBP2 knockdown cells. The graph shows the number of mKate2-FM4-PAUF-containing CARTS in the indicated cells at 15 min after the temperature shift to 37°C. Data are means ± s.e.m. (n = 11 cells per conditions, \**P* < 0.05, \*\*\*\**P* < 0.0001, one-way ANOVA multiple comparison test). Scale bar, 10 μm.

## Discussion

SCAP was discovered in 1996 as an ER protein whose mutation conferred, on CHO cells, resistance to 25-HC, an oxygenated cholesterol derivative that suppresses SREBP processing and thereby blocks cholesterol synthesis, but cannot replace cholesterol for cell viability^15^. Posterior studies demonstrated that SCAP senses the ER cholesterol content and, in response to cholesterol deficiency, escorts SREBPs from the ER to the Golgi complex for their cleavage-mediated activation^16–20^.

In the present study, we demonstrated a new function of SCAP under cholesterol-fed conditions. Based on our data, we propose a model whereby SCAP interacts with VAP/OSBP at ER-Golgi contact sites via Sac1 and regulates counter-transport of cholesterol and PI4P in an ER cholesterol-dependent manner (Fig. 9). This idea is supported by our finding that SCAP knockdown causes accumulation of PI4P at the TGN (Fig. 4b,c), a hallmark of the impairment of cholesterol/PI4P exchange between the ER and the *trans*-Golgi membranes. SCAP appears to contribute to the efficient establishment of ER-Golgi contact sites because its knockdown caused partial redistribution of the VAP/OSBP complex to ER-endosome contact sites (Fig. 4d,e). At the *trans*-Golgi membranes, cholesterol and SM organize lipid nanodomains, which can function as a platform for molecular machineries responsible for processing and sorting of cargoes, including GPI-anchored proteins (Fig. 9, right panel). In parallel, DAG, which is synthesized together with SM from ceramide and phosphatidylcholine, recruits PKD for membrane fission, leading to CARTS biogenesis. Intriguingly, our finding of putative nascent CARTS in the close vicinity to ER-Golgi contact sites (Fig. 5a) suggests a possible role of the ER contacts in determining the position for membrane fission, analogous to their important role in mitochondrial and endosomal fission^65,66^.

**Fig. 9.**
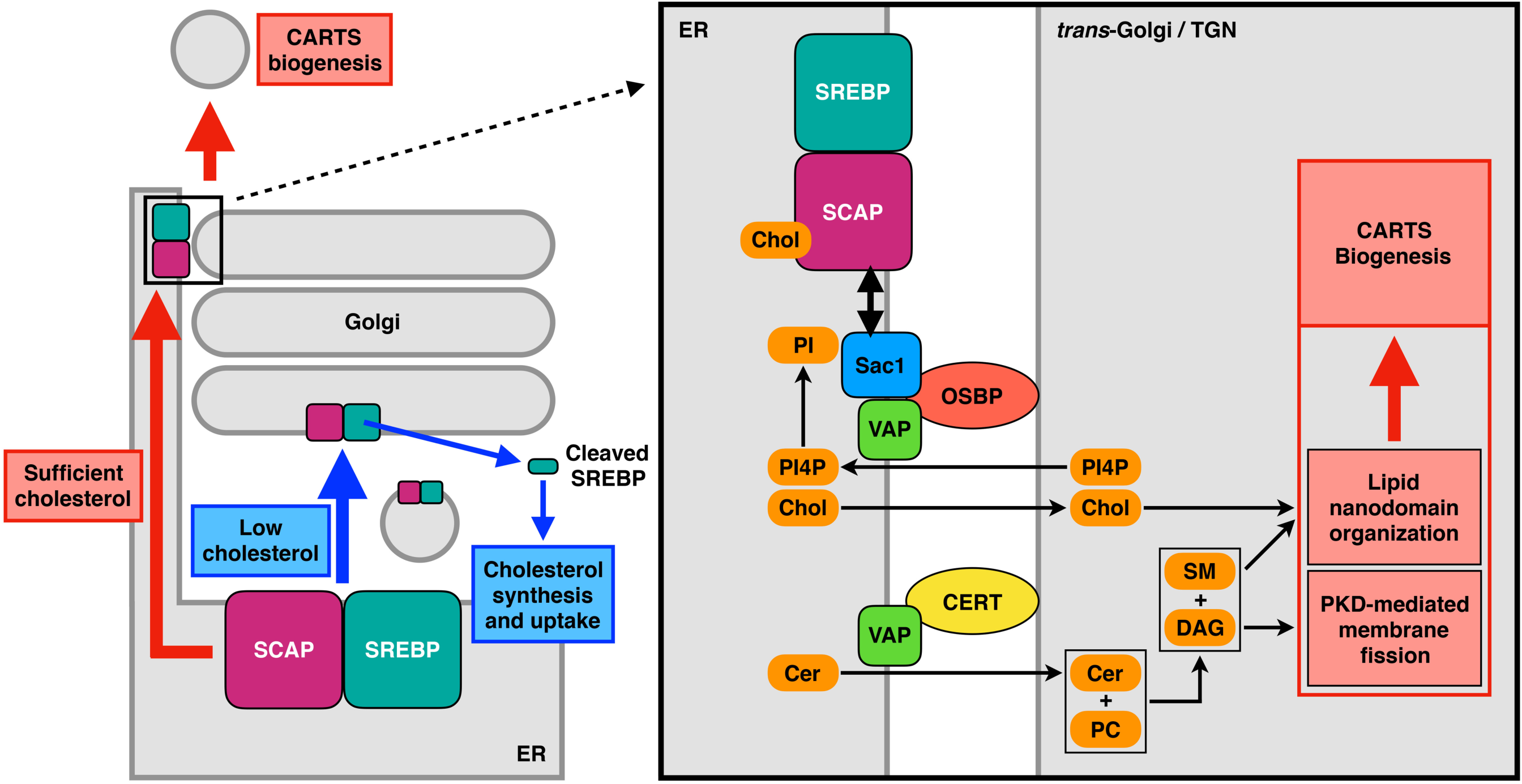
Working model for the facilitation of CARTS biogenesis by the SCAP/SREBP complex at ER-Golgi contact sites. When the ER cholesterol (Chol) level is low, SCAP escorts SREBP transcription factors from the ER to the Golgi complex for cholesterol synthesis and uptake (left panel, blue arrows). When the ER contains sufficient levels of cholesterol, a complex of cholesterol-bound SCAP and SREBP interacts with the VAP/OSBP complex via Sac1, and functions in the counter-transport of ER cholesterol and Golgi PI4P at ER-Golgi contact sites to promote CARTS biogenesis at the TGN domains immediately adjacent to the ER contacts sites (left panel, red arrows, and right panel). At these TGN domains, ceramide (Cer) transported by the VAP/CERT complex is metabolized together with phosphatidylcholine (PC) to SM and DAG. SM assembles with cholesterol into lipid nanodomains for processing and sorting of cargoes, while DAG recruits PKD for membrane fission, leading to CARTS biogenesis.

Cytoplasmic coats generally provide a means to coordinate signal-mediated cargo sorting with carrier budding and membrane fission. However, most TGN-derived transport carriers, including CARTS, lack cytoplasmic coats and therefore such molecular mechanisms remain largely elusive for these transport carriers^35,67^. A recent paper reported that Ca^2+^-dependent and oligomerization-driven cargo sorting, which is mediated by the SPCA1 Ca^2+^ pump and the secreted Ca^2+^ binding protein Cab45, is coupled to local SM synthesis at the TGN^14^. The finding that sorting of lysozyme C, one of the secretory cargoes in CARTS^35^, is controlled by this mechanism strongly supports our model.

Our data showed that neither the SCAP Y234A nor Y298C mutants, which reflect different conformations in the context of Insig binding, are competent for PI4P turnover and CARTS biogenesis (Fig. 7c,d). These findings imply that conformational switching that reflects cholesterol-free and -bound states is important. Considering that the ER contains low levels of cholesterol, it is tempting to speculate that SCAP functions in collecting cholesterol in the ER membrane to present it to cytosolic OSBP for efficient cholesterol transfer at ER-Golgi contact sites. It is of note that SCAP knockdown dramatically altered the distribution of the VAP/OSBP complex in the presence of 25-HC (Fig. 4d). As 25-HC directly binds to OSBP^41,42^, but indirectly to SCAP via Insig^68,69^, this finding reveals the possibility that SCAP functions at ER-Golgi contact sites as a complex with Insig, as well as SREBPs, supported by a previous finding that Insig interacts with VAP-A and VAP-B^70^.

In conclusion, our findings reveal a new role of SCAP under cholesterol-fed conditions and provide insights into the regulatory mechanisms for lipid transfer at ER-Golgi contact sites for transport carrier biogenesis at the TGN.

## Materials and methods

### Antibodies and reagents

Monoclonal antibodies were procured as follows: FLAG and alpha-tubulin were purchased from Sigma-Aldrich; Myc, hamster SCAP (clone: 9D5), and HMGR from Santa Cruz Biotechnology; SREBP1 from EMD Millipore; Penta-His from QIAGEN; CD63 (clone H5C6) from the Developmental Studies Hybridoma Bank; and PI4P from Echelon Biosciences. Polyclonal antibodies were procured as follows: HA, Nogo (RTN-4B), SCAP, and GST from Santa Cruz Biotechnology; TGN46 (sheep IgG) from AbD Serotec; GFP from Thermo Fisher Scientific; SREBP2, LDLR, and Golgin-97 from Abcam; and Myc from Cell Signaling Technology. To raise rabbit polyclonal antibodies against TGN46 and Sac1, GST-tagged fragments of human TGN46 (aa 1-365) and Sac1 (aa 1-55), respectively, were expressed in *Escherichia coli*, purified, and used as antigens. These antibodies were isolated by affinity chromatography on antigen-coupled beads. An anti-VAP-A and anti-Bap31 polyclonal antibodies were produced as described previously^34,39^. An anti-Mannosidase II polyclonal antibody was provided by V. Malhotra (Centre for Genomic Regulation, Barcelona, Spain). D/D solubilizer were purchased from Clontech. Methyl-β-cyclodextrin and 2-hydroxypropyl-β-cyclodextrin were purchased from Wako Chemicals. Filipin and 25-HC were purchased from Sigma-Aldrich.

### Plasmids

The plasmids encoding PAUF-MycHis, N-acetylglucosaminyl transferase I-GFP, Myc-OSBP^71^, and HA-CERT^72^ were kindly donated by S. S. Koh (Korea Research Institute of Bioscience and Biotechnology, Daejeon, Korea), N. Nakamura (Kyoto Sangyo University, Kyoto, Japan), H. Arai (University of Tokyo, Tokyo, Japan), K. Hanada (National Institute of Infectious Diseases, Tokyo, Japan), respectively. The plasmids encoding mKate2-FM4-GPI and EQ-SM (tagged with oxGFP) were generous gifts from C. G. Burd (Yale School of Medicine, New Haven, CT, USA). The plasmids encoding the GFP-Sac1 WT and K2A mutant were generous gifts from P. Mayinger (Oregon Health and Science University, Portland, OR, USA). The plasmids encoding FLAG-SREBP1a and FLAG-SREBP2 were generous gifts from D. M. Sabatini (Whitehead Institute for Biomedical Research, Cambridge, MA, USA) (Addgene plasmids #32017 and #32018)^73^. The plasmid encoding GST-PKD2 kinase dead was provided by V. Malhotra (Centre for Genomic Regulation, Barcelona, Spain). The cDNAs encoding hamster SCAP WT and C-term (aa 732-1276) were amplified by PCR from pCMV-GFP-SCAP, a generous gift from P. Espenshade (Johns Hopkins University School of Medicine, Baltimore, MD, USA), and inserted into pFLAG-CMV-6c to express the protein with an N-terminal FLAG. For expression of FLAG-SCAP D451A/L452A, point mutations were introduced into pFLAG-SCAP by PCR using primers designed for replacing aspartic acid 451 and leucine 452 with alanines. The cDNA encoding hamster SCAP or rabbit OSBP (WT or FF/AA) were inserted into pCI-IRES-Bsr in combination with one encoding the N-terminal fragment of Venus (Vn) (aa 1-172). The cDNA encoding human Sac1 or VAP-A was inserted into pCI-IRES-Bsr in combination with one encoding the C-terminal fragment of Venus (Vc) (aa 154-238). For establishment of stable cell lines, the cDNAs encoding Vn-OSBP and Vc-VAP-A were inserted into the pMXs-IRES-Puro and pCX4-IRES-Bsr retroviral vectors, respectively, and hamster SCAP with or without N-terminal FLAG or GFP was inserted into pCX4-IRES-Bsr. For expression of SCAP Y234A or Y298C, each point mutation was introduced into pCX4-SCAP-IRES-Bsr by PCR using primers designed for replacing tyrosine 234 or 298 with alanine. For expression of mKate2-FM4-PAUF, the cDNA encoding GPI in the plasmid for mKate2-FM4-GPI was replaced with one encoding human PAUF (aa 43-197). The cDNA encoding human sialyltransferase (aa 1-69) was inserted into a pcDNA3-based plasmid encoding mRFP to express the protein with a C-terminal mRFP. Plasmids for Myc-OSBP FF/AA and PH-FFAT were described previously^34^. For establishment of shSCAP HeLa cells, the Mission shRNA plasmid (TRCN0000289279) was purchased from Sigma-Aldrich.

### siRNA and shRNA

The targeting sequences of siRNA and shRNA were as follows:

Control (GL2 luciferase): 5’-AACGTACGCGGAATACTTCGA-3’

SCAP (siRNA): 5’-AACCTCCTGGCAGTAGATGTA-3’

SCAP (shRNA): 5’- GCTCTGGTGTTCTTGGACAAA-3’

VAP-A: 5’-AACTAATGGAAGAGTGTAAAA-3’

VAP-B: 5’-AAGAAGGTTATGGAAGAATGT-3’

OSBP: 5’-AATACTGGGAGTGTAAAGAAA-3’

CERT: 5’-AAGAACAGAGGAAGCATATAA-3’

Sac1: 5’-AACTGATATTCAGTTACAAGA-3’

SREBP1: 5’-GAGGCAAAGCTGAATAAATCT-3’

SREBP2: 5’-CAGGCTTTGAAGACGAAGCTA-3’

### Cell culture and transfection

HeLa and HEK 293T cells were grown in DMEM supplemented with 10% FCS. PLAT-A packaging cells were grown in DMEM supplemented with 10% FCS, 10 μg/ml blasticidin S, and 1 μg/ml puromycin. Plasmid and siRNA transfection into HeLa cells were carried out using X-tremeGENE 9 DNA transfection reagent (Roche) and Oligofectamine reagent (Invitrogen), respectively, according to the manufacturers’ protocols. Plasmid transfection into HEK 293T cells was carried out using polyethylenimine (Polysciences) or Lipofectamine 2000 transfection reagent (Invitrogen), according to the manufacturers’ protocols.

### Establishment of stable cell lines

For establishment of HeLa cells stably expressing FLAG- or GFP-SCAP, PLAT-A packaging cells were transfected with pCX4-FLAG- or GFP-SCAP-IRES-Bsr, and 48 h later, the medium containing retrovirus was collected and used for infection of HeLa cells. Selection of HeLa cells stably expressing FLAG- or GFP-SCAP was performed with 10 μg/ml blasticidin S. A HeLa stable cell line coexpressing Vn-OSBP and Vc-VAP-A was established in a similar manner with pMXs-Vn-OSBP-IRES-Puro and pCX4-Vc-VAP-A-IRES-Bsr by selecting cells with 1 μg/ml puromycin and 10 μg/ml blasticidin S. For establishment of shSCAP HeLa cells, HEK 293T cells were cotransfected with the Mission shRNA plasmid (TRCN0000289279), pMDLg/pRRE, pRSV-Rev, and pCMV-VSVG. After 48 h, the medium containing lentivirus was collected and used for infection of HeLa cells. Selection of shSCAP HeLa cells was performed with 1 μg/ml puromycin. Establishment of shSCAP HeLa cells stably expressing hamster SCAP WT, Y234A, or Y298C was performed as described for GFP-SCAP HeLa cells with pCX4-SCAP (WT, Y234A, or Y298C)-IRES-Bsr. All the stable cell lines were obtained without single-cell cloning.

### Immunofluorescence microscopy

HeLa cells were fixed with 4% paraformaldehyde (PFA) in phosphate-buffered saline (PBS) at room temperature for 20 min, permeabilized with 0.2% Triton X-100 in PBS for 30 min, and then blocked with 2% bovine serum albumin (BSA) in PBS for 30 min. The cells were labeled with the indicated primary antibodies and secondary antibodies conjugated to Alexa Fluor 488, 594, or 633 in the blocking buffer. The samples were analyzed with an Olympus Fluoview FV1000 or FV1200 confocal microscope with a UPLSAPO 60x O NA 1.35 objective and FV10-ASW software. Image processing and measurement of fluorescence intensity were performed with ImageJ software (National Institutes of Health). Vesicular structures containing EQ-SM and/or PAUF-MycHis were detected using the plugin ComDet (https://github.com/ekatrukha/ComDet).

### Image deconvolution

Deconvolution processing was performed with Huygens Professional version 18.04 (Scientific Volume Imaging, The Netherlands, http://svi.nl). For that, a theoretical point-spread function (PSF) was automatically computed based on the microscope and image acquisition parameters. The deconvolution process was numerically performed using the Classic Maximum Likelihood Estimation (CMLE) algorithm. In brief, this algorithm assumes that the photon noise is Poisson-distributed, and the likelihood of an estimate of the actual image given the computed PSF and the acquired image is iteratively optimized until either a quality factor or a maximum number of iterations is reached. In our deconvolutions, we used a quality factor equal to 0.001 and a maximum 50 iterations. The signal-to-noise ratio (SNR) for each acquired image was computed based on three line profiles going through regions of background signals towards regions of positive, actual signals. Typically, SNR were of the order of 7–20 for the different analyzed images.

### Filipin staining

Filipin staining was performed as described previously^74^. In brief, HeLa cells were treated with 10 mM Methyl-β-cyclodextrin at 37°C for 30 min and fixed with 4% PFA at room temperature for 10 min. The cells were incubated with 0.1 mg/ml filipin in PBS at room temperature for 30 min. After washing with PBS, the cells were blocked with 1% BSA in PBS for 30 min and labeled with anti-Golgin-97 antibody and a secondary antibody conjugated to Alexa Fluor 594 in the blocking buffer. After re-incubation with 0.1 mg/ml filipin at room temperature for 30 min, the cells were washed with PBS and then mounted on a microscope slide. The samples were analyzed by fluorescence microscopy as described above.

### PI4P staining

PI4P staining was performed as described previously^13,75^. In brief, HeLa cells were fixed with 2% PFA at room temperature for 15 min, followed by washing with PBS containing 50 mM NH_4_Cl. The cells were then permeabilized with 20 μM digitonin in buffer A (20 mM PIPES, pH 6.8, 137 mM NaCl, 2.7 mM KCl). After removal of digitonin by washing with buffer A, the cells were blocked with 5% FCS in buffer A for 45 min. The cells were labeled with anti-PI4P and anti-Golgin-97 antibodies, followed by secondary antibodies conjugated to Alexa Fluor 488 and 594, respectively, in the blocking buffer. After post-fixation with 2% PFA for 5 min, the cells were washed with PBS containing 50 mM NH_4_Cl and with water, and then mounted on a microscope slide. The samples were analyzed by fluorescence microscopy as described above.

### Quantification of Golgi PI4P

To quantify the intensity of the PI4P signal at the Golgi complex from our immunofluorescence microscopy images, we used the ImageJ distribution Fiji^76^ and a custom-made macro, following the method described in Ref^77^. First, we generated a binary mask for the Golgi/TGN area using the signal of the marker protein Golgin-97 after 1-pixel radius Gaussian blur filtering. Then, the mean intensities of the PI4P and Golgin-97 channels in the Golgi/TGN mask areas were background subtracted and measured separately for each individual cell. Finally, the measured Golgi (mask) PI4P levels were normalized to the Golgi (mask) Golgin-97 levels per each cell.

### Quantification of perinuclear BiFC signal

To quantify the perinuclear BiFC signal from our fluorescence microscopy images, we used the ImageJ distribution Fiji and a custom-made macro. First, we generated a binary mask for the perinuclear area using the signal of the TGN marker protein TGN46 after 1-pixel radius Gaussian blur filtering. Finally, the total intensity of the BiFC channel in the perinuclear mask area was measured separately for each individual cell after background subtraction.

### iFRAP

HeLa cells expressing GFP-Sac1 WT, K2A, or N-acetylglucosaminyl transferase I-GFP in Opti-MEM were cultured in 5% CO_2_ at 37°C during live-cell imaging. The cells were subjected to bleaching with high laser intensity (473 nm laser) for 15 s, followed by an imaging scan with a time interval between frames of 10 s for ∼4 min by use of an Olympus Fluoview FV1000 confocal microscope with a UPLSAPO 100x O NA 1.40 objective and FV10-ASW software. Image processing and measurement of fluorescence intensity were performed with ImageJ software.

### Immunoprecipitation

HEK 293T cells were lysed in buffer B (50 mM HEPES-KOH, pH 7.4, 100 mM NaCl, 1.5 mM MgCl_2_, 1 mM dithiothreitol, 1% Nonidet P-40, 1 μg/ml leupeptin, 2 μM pepstatin A, 2 μg/ml aprotinin, and 1 mM phenylmethylsulfonyl fluoride). The lysates were centrifuged at 17,000 × g for 10 min. The resulting supernatants were immunoprecipitated with an anti-FLAG M2 affinity gel (Sigma-Aldrich), and the precipitated proteins were analyzed by Western blotting with the indicated primary antibodies and secondary antibodies conjugated to horseradish peroxidase. For immunoprecipitation of FLAG-SREBPs and stably expressed FLAG-SCAP, cells were lysed in buffer B without dithiothreitol and the lysates were centrifuged at 17,000 × g for 10 min. The supernatants were further centrifuged at 100,000 × g for 30 min and the resulting supernatants were immunoprecipitated as described above. For identification of the Sac1 interacting proteins, the precipitated proteins were eluted with FLAG peptide and analyzed by silver staining. Protein bands were excised from the gel and subjected to LC-MS/MS analysis.

### LC-MS/MS analysis

Each gel piece was incubated with 0.25 μg trypsin in 20 μl Tris-HCl (pH 8.8) overnight at 37°C^78^, and the resulting peptide mixture was analyzed with a direct nanoflow LC-MS/MS system equipped with an hybrid quadrupole-orbitrap mass spectrometer (Q Exactive, Thermo Scientific, Boston, MA) as previously described^79^. Briefly, peptides were separated on a reversed-phase tip column (150 μm i.d. × 70 mm, Mightysil-C18, 3-μm particle) using a 0–35% linear gradient of acetonitrile in 0.1% (v/v) formic acid for 35 or 70 min at a flow rate of 100 nL/min. Full MS scans were acquired with a resolution of 30,000 at a mass-to-charge ratio of 400. The ten most intense ions were fragmented in the data-dependent mode by collision-induced dissociation with normalized collision energy of 35, activation q of 0.25, and activation time of 10 ms and one microscan. The MS/MS data were converted to the mascot generic format with the Proteome Discoverer software (Thermo Scientific, ver. 1.1). The files were processed with the MASCOT algorithm (version 2.2.1., Matrix Science Ltd., London, United Kingdom) to assign peptides using the Swiss-Prot sequence database (release 2012.11, human) under the search parameters described in Ref^79^. Peptides were identified based on the MASCOT definitions. For the search parameters, we set the variable modifications for acetylation (protein N terminus) and oxidation (Met). The maximum missed cleavage was set at 1 with a peptide mass tolerance of +/-15 ppm. Peptide charges from +2 to +4 states and MS/MS tolerances of +/-0.8 Da were allowed. All results of peptide searches were extracted from the Mascot DAT files using the STEM software^80^.

### Cholesterol depletion

Lipoprotein-deficient serum (LPDS) was prepared as described previously^81,82^. In brief, FCS was adjusted to a density of 1.25 g/mL with solid KBr and then centrifuged for 16 h at 10°C at 50,000 rpm in a Beckman VTi 50 rotor. The upper yellow-orange layer containing lipoproteins and the salt pellet were removed, and the remaining fraction was dialyzed extensively at 4°C against 150 mM NaCl for 48 h. HeLa cells were incubated with DMEM containing 5% LPDS and 1% 2-hydroxypropyl-β-cyclodextrin for 3 h.

### Quantitative real-time PCR

RNA was prepared using an RNeasy Mini kit (QIAGEN) and cDNA was synthesized with SuperScript III reverse transcriptase (Invitrogen) primed by oligo(dT)15. Quantitative real-time PCR was performed with a Rotor-Gene Q RT-PCR machine (QIAGEN) using a KAPA SYBR FAST qPCR kit (Kapa Biosystems), according to the manufacturer’s protocol. The primers used were as follows: SCAP, forward primer 5’- TATCTCGGGCCTTCTACAACC-3’ and reverse primer 5’- GGGGCGAGTAATCCTTCACA-3’; HMGR, forward primer 5’- TGACCTTTCCAGAGCAAGC-3’ and reverse primer 5’- CCAACTCCAATCACAAGACATTC-3’; LDLR, forward primer 5’- GTGTCACAGCGGCGAATG-3’ and reverse primer 5’- CGCACTCTTTGATGGGTTCA-3’; HPRT1, forward primer 5’- TTCCAGACAAGTTTGTTGTAGGAT-3’ and reverse primer 5’- GCAGATGGCCACAGAACTAG-3’. Values were normalized as the HPRT1 expression level.

### Total cholesterol measurement

HeLa cells were lysed in water for 30 min at 37°C and then subjected to total cholesterol measurement with an Amplex Red cholesterol assay kit (Molecular probes), according to the manufacturer’s protocol. Values were normalized to protein content determined as using a Pierce BCA protein assay kit (Thermo Fisher Scientific).

### PAUF secretion assay

HeLa cells were transfected with control siRNA or siRNA oligos targeting SCAP. At 48 h after siRNA transfection, the cells were transfected with a plasmid for PAUF-MycHis, 20 h later the medium was replaced with Opti-MEM, and then the cells were incubated at 37°C for 6 h. After collecting the medium, the cells were lysed with 0.5% sodium dodecyl sulfate (SDS) and 0.025 units/μL benzonase nuclease (Sigma-Aldrich) in PBS. The medium and cell lysates were analyzed by Western blotting with an anti-Penta-His antibody.

### GPI transport assay

HeLa cells were transfected with control siRNA or siRNA oligos targeting SCAP, VAP-A/VAP-B, OSBP or CERT/OSBP. At 48 h after siRNA transfection, the cells were transfected with a plasmid for mKate2-FM4-GPI, 20 h later the medium was replaced with DMEM containing 10% FCS, 1 μM D/D solubilizer, and 20 μg/mL cycloheximide, and then cells were incubated at 37°C for the indicated times. The cells were then fixed with 4% PFA and analyzed by fluorescence microscopy as described above.

### Inducible CARTS formation assay

HeLa cells were transfected with control siRNA or siRNA oligos targeting SCAP, SREBP1, or SREBP2. At 48 h after siRNA transfection, the cells were transfected with a plasmid for mKate2-FM4-PAUF, 20 h later the medium was replaced with DMEM containing 10% FCS, 20 mM HEPES-KOH, pH 7.4, 1 μM D/D solubilizer, and 20 μg/mL cycloheximide, and then the cells were incubated in a water bath at 20°C for 45 min. The cells were then incubated in a water bath at 37°C for the indicated times, followed by fixation with 4% PFA. The samples were analyzed by fluorescence microscopy as described above. Quantification of CARTS was performed with ImageJ software (National Institutes of Health). The fluorescence signal of mKate2-FM4-PAUF-containing puncta was distinguished from the background by setting a threshold and analyzed at a set size within 0.05-2.00 μm^2^. For live-cell imaging, HeLa cells were transfected with a plasmid for mKate2-FM4-PAUF, 20 h later the medium was replaced with Opti-MEM containing 1 μM D/D solubilizer and 20 μg/mL cycloheximide, and then the cells were incubated in a water bath at 20°C for 45 min. The cells were then incubated at 37°C and 5% CO_2_ to monitor CARTS biogenesis. Images were acquired continuously with a time interval between frames of 30 sec for ∼75 min by use of an Olympus Fluoview FV1200 confocal microscope with a UPLSAPO 60x O NA 1.35 objective and FV10-ASW software. The images were processed with ImageJ software.

## Acknowledgements

We thank Peter Espenshade, Peter Mayinger, Hiroyuki Arai, Kentaro Hanada, Nobuhiro Nakamura, Sang Seok Koh, David M. Sabatini, Cristopher G. Burd, and Vivek Malhotra for providing materials. We appreciate the technical assistance of Nanako Oyama, So Yoshida, Yuika Komatsuda, Yuiko Kawai, Natsumi Hoshino, Tomoya Iizuka, Sho Furuichi, and Katsunori Iwasa. We are grateful to Josse van Galen and Vivek Malhotra for comments on the manuscript. This work was supported in part by Grants-in-Aid for Scientific Research [grant numbers 15K18507 and 17K07348 to Y.W., and 18H02439 to M.T.] from the Ministry of Education, Culture, Sports, Science, and Technology of Japan, the Naito Foundation [to Y.W.], and the Ono Medical Research Foundation [to Y.W.]. F.C. acknowledges financial support from the Spanish Ministry of Economy and Competitiveness (“Severo Ochoa” program for Centres of Excellence in R&D (SEV-2015-0522), FIS2015 - 63550 - R, FIS2017 - 89560 - R, BFU2015-73288-JIN, AEI/FEDER/UE, and RYC-2017-22227), Fundació Privada Cellex, and from the Generalitat de Catalunya through the CERCA program.

## Author contributions

Y.W. and M.Tagaya conceived and designed the experiments. Y.W., K.H., T.N., C.W., F.C., and H.K. performed the experiments. M.Taoka performed LC-MS/MS analysis and analyzed the data. T.U., H.I., K.A., and M.Tagaya provided samples. Y.W., F.C., and M.Tagaya wrote the manuscript.

## Figure legends

**Supplementary Fig. 1.**
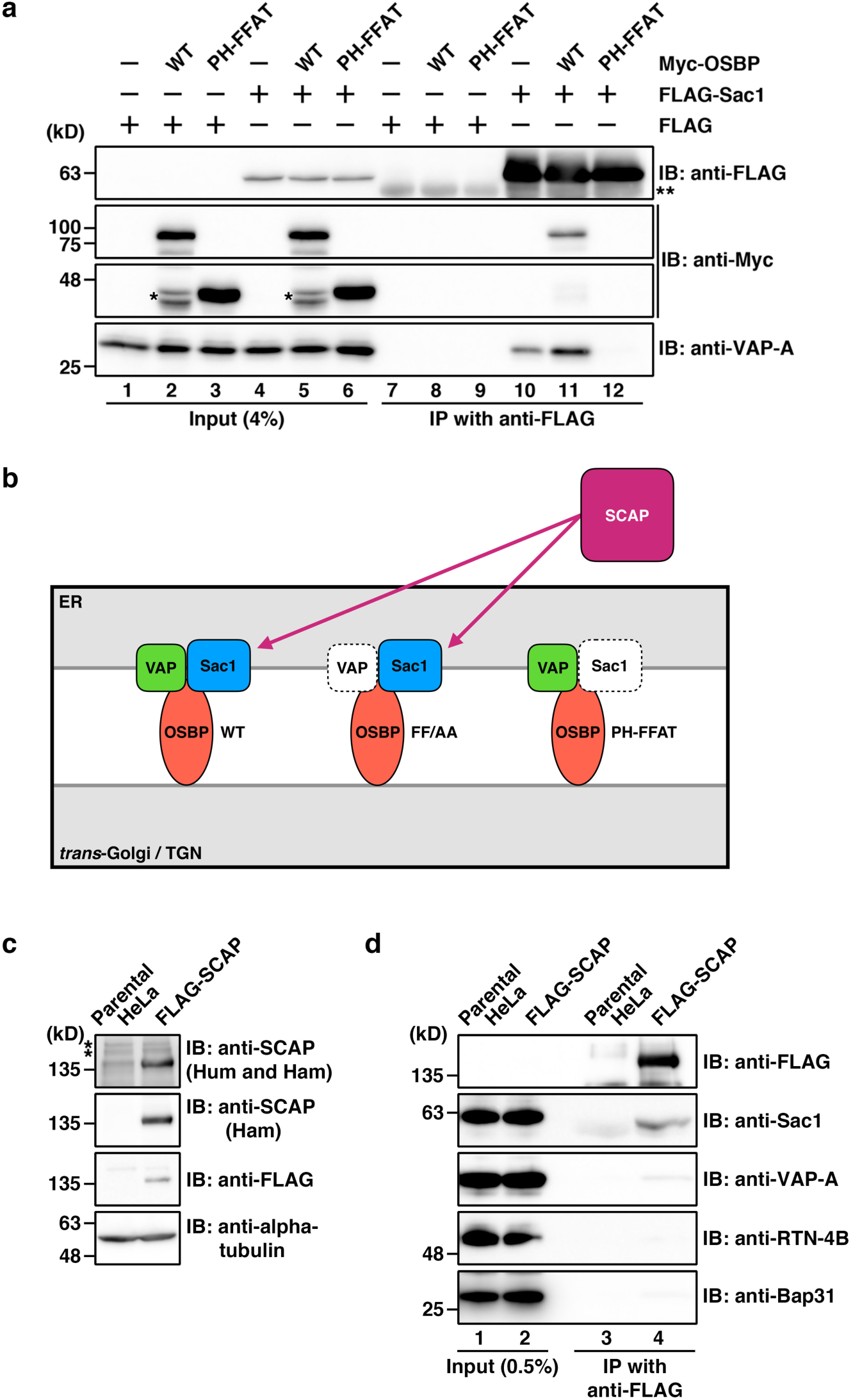
Characterization of the interaction of FLAG-Sac1 with Myc-OSBP PH-FFAT mutant, and the interaction of stably-expressed FLAG-SCAP with Sac1. **a**, Interactions of FLAG-Sac1 with VAP-A and Myc-OSBP WT, but not with the PH-FFAT mutant in HEK 293T cells. Cell lysates were subjected to immunoprecipitation with an anti-FLAG M2 affinity gel, and the cell lysates (Input) and immunoprecipitates (IP) were immunoblotted (IB) with the indicated antibodies. The interaction of FLAG-Sac1 with VAP-A was enhanced by coexpression with Myc-OSBP WT, but not with PH-FFAT. Single and double asterisks denote degraded Myc-OSBP fragments and the immunoglobulin heavy chain, respectively. **b**, A schematic representation of the SCAP interactions with VAP, OSBP, and Sac1. **c**, Establishment of a HeLa stable cell line expressing FLAG-SCAP. Cell lysates were immunoblotted with indicated antibodies. One of two SCAP antibodies recognizes both human (Hum) and hamster (Ham) SCAP and the other is hamster-specific. Asterisks denote nonspecific bands. **d**, Interaction of stably-expressed FLAG-SCAP with Sac1, but not with RTN-4B and Bap31. VAP-A was detected as a very faint band in the immunoprecipitate of the stable cell line (lane 4).

**Supplementary Fig. 2.**
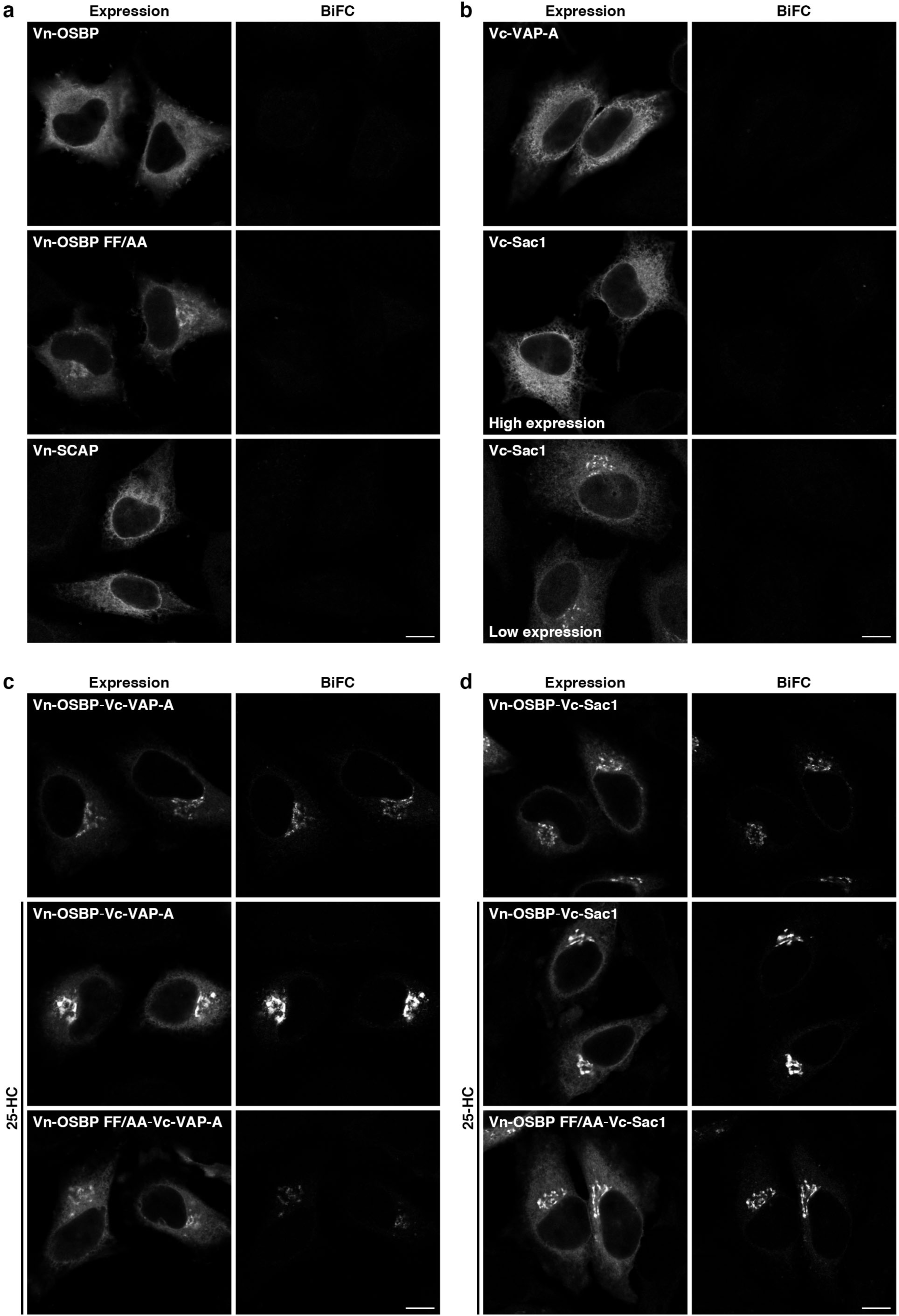
BiFC visualization of VAP-A, OSBP, and Sac1 interactions at ER-Golgi contact sites. **a**,**b**, No BiFC signal in HeLa cells with single expression of Venus N-terminal fragment (Vn)-fused proteins (**a**) or Venus C-terminal fragment (Vc)-fused proteins (**b**). **c**,**d**, BiFC visualization of OSBP/VAP-A (**c**) and OSBP/Sac1 (**d**) interactions at ER-Golgi contact sites. The coexpression of Vn-OSBP (WT or FF/AA) with Vc-VAP-A (**c**) or Vc-Sac1 (**d**) was visualized with an anti-GFP antibody. The BiFC signal was enhanced by treatment of cells with 2 μg/mL 25-HC for 2.5 h. Vn-OSBP FF/AA showed a reduced BiFC signal, compared with the WT. Scale bars, 10 μm.

**Supplementary Fig. 3.**
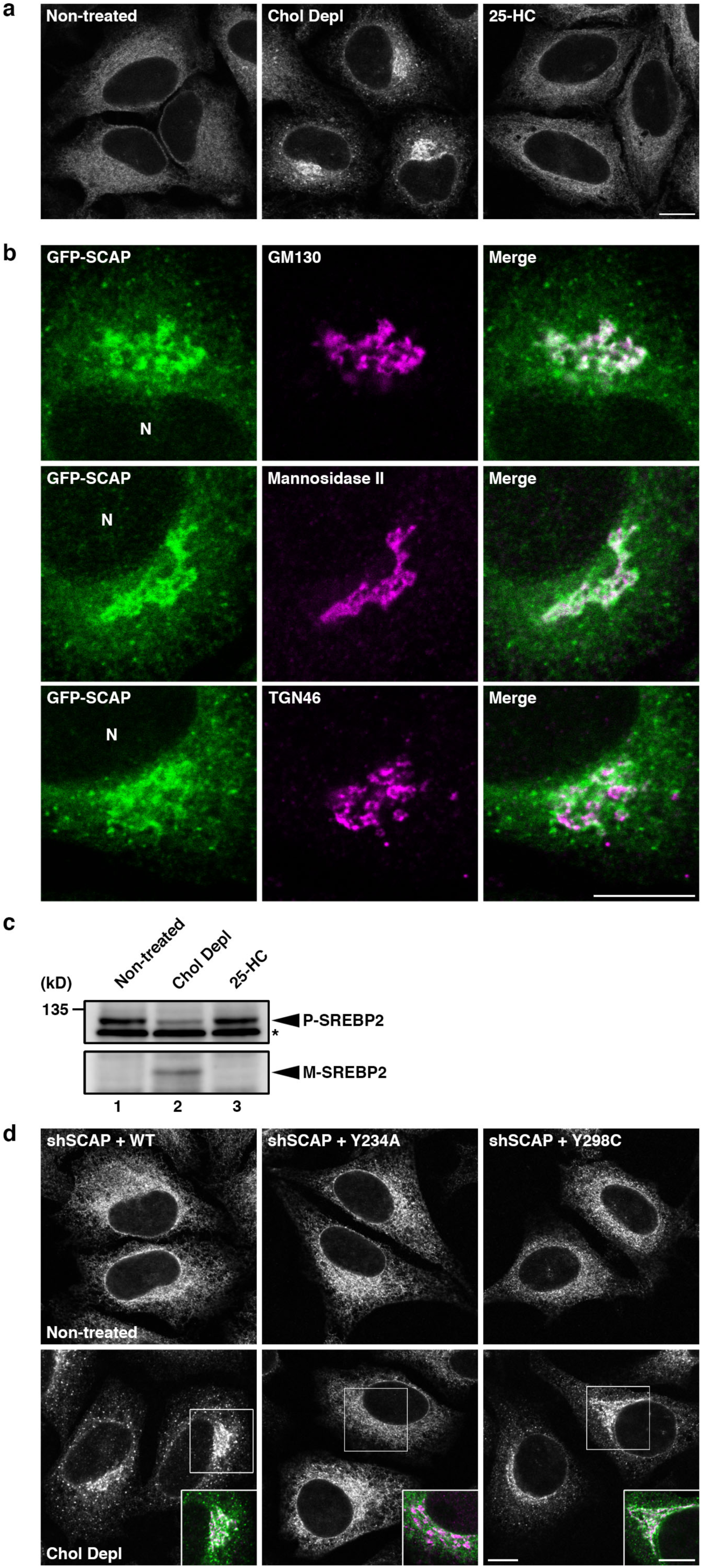
Sterol-dependent localization of SCAP. **a**,**b**, Localization of stably expressed GFP-SCAP in HeLa cells with or without cholesterol depletion (Chol Depl) for 3 h or treated with 2 μg/mL 25-HC for 2.5 h. Accumulation of GFP-SCAP in GM130 (*cis*-Golgi matrix protein)- and Mannosidase II (*cis*/*medial*-Golgi marker enzyme)-positive membranes upon cholesterol depletion for 3 h (**b**). N, nucleus. **c**, Proteolytic activation of SREBP2 upon cholesterol depletion, monitored by Western blotting with an anti-SREBP2 antibody. Arrowheads indicate the precursor (P) and mature (M) forms of SREBP2. Asterisk denotes nonspecific bands. **d**, Localization of the stably-expressed hamster SCAP WT, Y234A, and Y298C mutants in shSCAP HeLa cells with or without cholesterol depletion for 3 h. Merged images for the hamster SCAP WT, Y234A or Y298C (green) and the *cis*/*medial*-Golgi marker GPP130 (magenta) of the boxed areas are shown in the insets. Scale bars, 10 μm.

**Supplementary Fig. 4.**
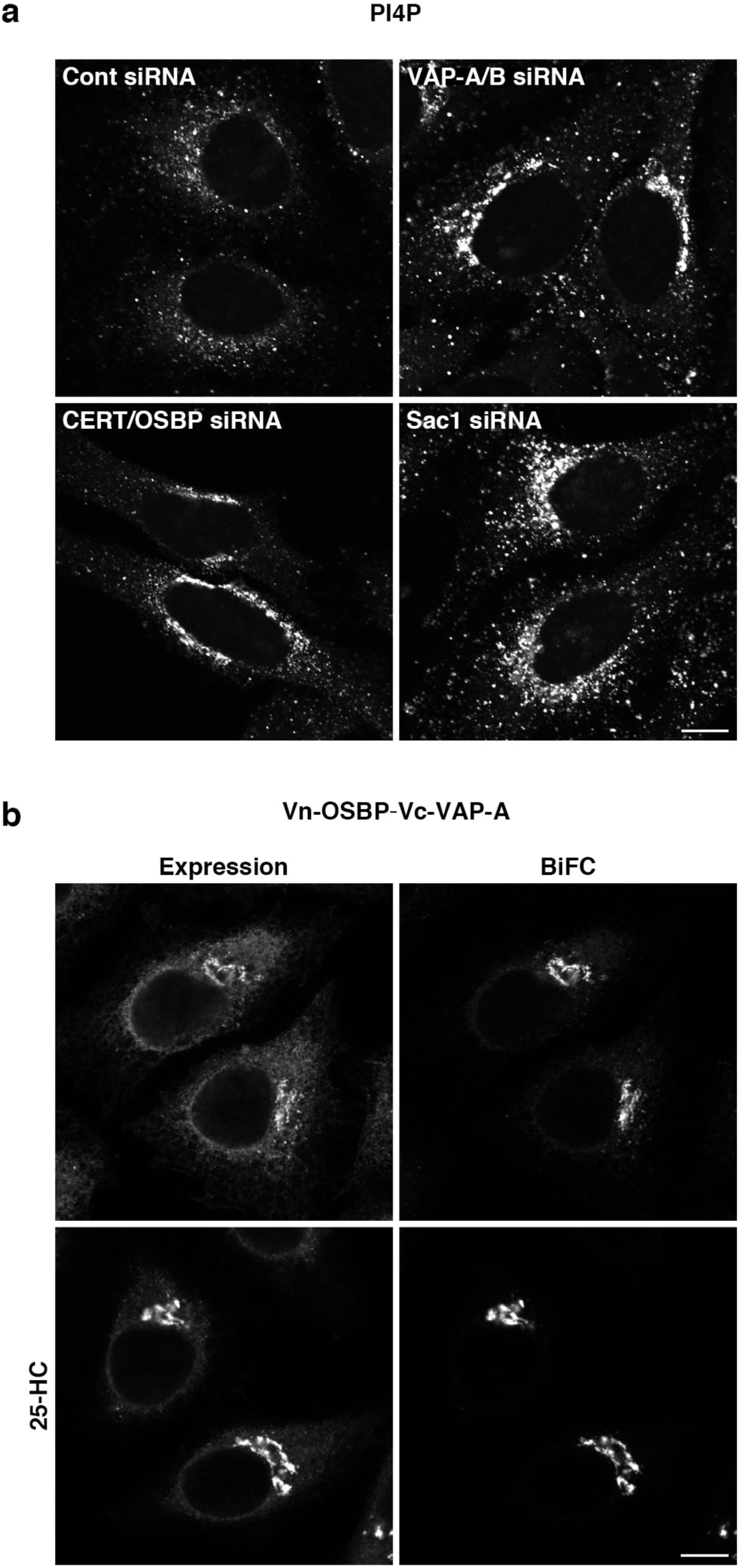
Effects of knockdown of ER-Golgi contact site components on PI4P turnover, and visualization of VAP-A/OSBP complex at ER-Golgi contact sites. **a**, PI4P staining in control (Cont), VAP-A/B, CERT/OSBP, and Sac1 knockdown HeLa cells. **b**, HeLa cells stably coexpressing Vn-OSBP and Vc-VAP-A were treated with or without 2 μg/mL 25-HC for 2.5 h. The coexpression of Vn-OSBP and Vc-VAP-A was visualized with an anti-GFP antibody. Scale bars, 10 μm.

**Supplementary Fig. 5.**
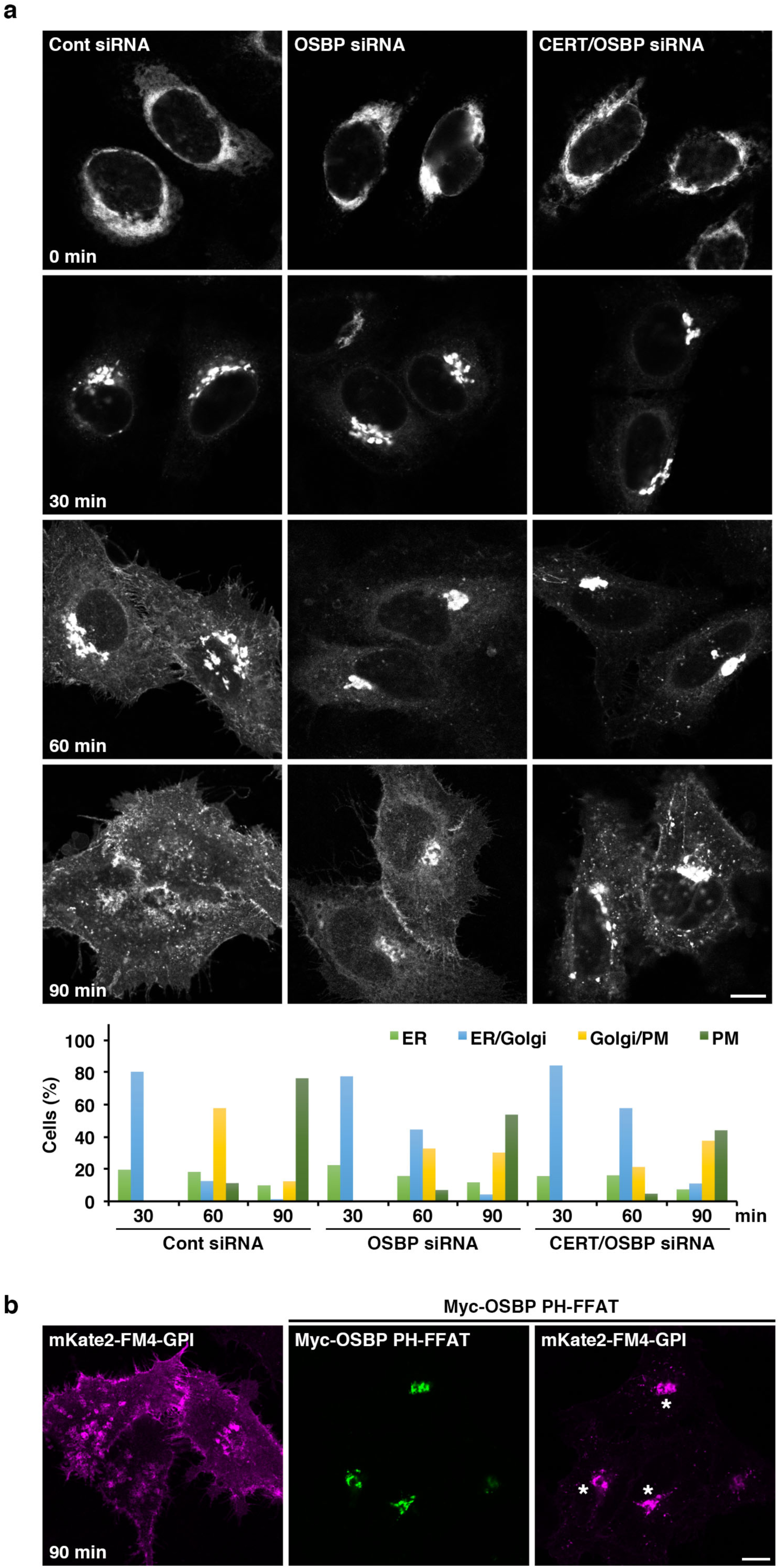
Lipid transfer complexes at ER-Golgi contact sites are required for GPI-anchored protein transport from the TGN to the PM. **a**, mKate2-FM4-GPI transport from the ER to the PM via the Golgi complex in control (Cont), OSBP, and CERT/OSBP knockdown cells. The cells were incubated at 37°C with the D/D solubilizer and cycloheximide, and fixed at the indicated times. The graph shows the percentages of cells with mKate2-FM4-GPI at the ER, ER/ Golgi, Golgi/PM, or PM at the indicated times. The data shown are for a single representative experiment out of three performed (control siRNA: 30 min, n = 240 cells; 60 min, n = 230 cells; 90 min, n = 233 cells; OSBP siRNA: 30 min, n = 254 cells; 60 min, n = 229 cells; 90 min, n = 238 cells; CERT/OSBP siRNA: 30 min, n = 268 cells; 60 min, n = 235 cells; 90 min, n = 245 cells). **b**, mKate2-FM4-GPI transport in the presence or absence of the Myc-OSBP PH-FFAT mutant. Asterisks denote cells coexpressing mKate2-FM4-GPI and Myc-OSBP PH-FFAT. Scale bars, 10 μm.

**Supplementary Video 1 | The biogenesis of mKate2-FM4-PAUF-containing CARTS at the TGN.** HeLa cells expressing mKate2-FM4-PAUF were first incubated at 20°C with the D/D solubilizer and cycloheximide for 45 min. Images were acquired continuously after the temperature shift to 37°C with a time interval between frames of 30 sec for ∼75 min. N, nucleus.

